# Self-Assembled Nucleolipid G-Quadruplexes Act as Multitarget Decoys for Oncogene Suppression in Pancreatic Cancer

**DOI:** 10.64898/2026.04.03.715535

**Authors:** Faith Kivunga, Virginie Baylot, Tina Kauss, Brune Vialet, Jules Simonin Garcia, Patricia Korczak, Zahraa Othman, Gilmar Salgado, Philippe Barthélémy

## Abstract

KRAS mutations drive multiple cancers and represent an important therapeutic target, together with other oncogenic regulators such as MYC, KIT, and BCL2 that are critically involved in pancreatic cancer. Here we describe a novel therapeutic strategy based on stable nucleolipid-modified G-quadruplexes (NLG4). Cell viability assays demonstrate that NLG4 strongly inhibit pancreatic cancer cell proliferation, whereas non-lipidic G-quadruplex sequences display minimal activity under comparable conditions. Owing to their distinctive physicochemical properties, including stabilization of parallel G-quadruplex structures and self-assembly into micellar aggregates, NLG4 efficiently internalize into cells and interact with key G-quadruplex unfolding factors such as UP1. This interaction leads to a marked downregulation of KRAS, c-MYC, c-KIT, and BCL2 expression. Suppression of these oncogenes profoundly affects pancreatic cancer cell fate, as evidenced by reduced expression of proliferation (Ki67) and anti-apoptotic (BCL2) markers. In addition, NLG4 treatment decreases inflammatory signaling mediated by NF-κB and inhibits major pro-proliferative kinase pathways, including ERK, AKT, and phosphorylated AKT. The therapeutic relevance of this decoy strategy is further supported by the observed potentiation of gemcitabine antitumor activity. Overall, these findings highlight NLG4 as a promising anticancer approach that simultaneously targets multiple oncogenic pathways through G-quadruplex-based decoy mechanisms, with translational potential for future pancreatic cancer treatment.

## 1. INTRODUCTION

Mutations in the *KRAS* gene are involved in different tumor types and due to its role in the development of cancer, this oncogene is an attractive target for new anticancer drugs and strategies. Indeed, KRAS is an important element in the pathogenesis of several type of cancers and a primary target for anticancer drugs ^1^. *KRAS* is the most frequently mutated oncogene in human cancer; *i.e.* mutated in more than 90% of pancreatic adenocarcinomas (PDAC) and in 50% of colorectal carcinomas, for example. If direct small-molecule inhibitors of KRAS are currently under investigation ^2,3^, new strategies are needed to specifically address this oncogenic target. MYC is one of the most common activated oncogene in human cancer, it is overexpressed in more than 40% of primary pancreatic cancers ^4,5,6^. Also, it was found that activation of MYC could convert pancreatic intraepithelial neoplasm into pancreatic cancer in mice ^7^. It was observed in preclinical studies that inhibition of c-*Myc* oncogene in resistant pancreatic tumors could improve the response to chemotherapies ^8^. These oncogenes are among the most frequently altered in cancers, with KRAS primarily mutated and MYC commonly amplified. Their cooperative interaction profoundly shapes tumor development, regulating key cancer hallmarks such as apoptosis evasion, limitless proliferation, angiogenesis, metastasis, metabolic reprogramming, and immune escape ^9^. Despite their central roles, both oncogenes were long deemed “undruggable”. Recent advances, however, have yielded promising inhibitors targeting KRAS ^10,11,12^ or MYC ^13,14^ now in clinical trials, or approved ^15^.

The interplay between MYC dysregulation and resistance to KRAS inhibition, but also the central role in cancer progression of oncogenic proteins such as tyrosine kinase KIT ^16,17^ and survival factor BCL2 ^18^, highlight the potential therapeutic benefit of targeting simultaneously these oncogenes. Demonstrating this synergy in preclinical models would be highly desirable and a critical step toward guiding therapeutic design and enhancing the likelihood of successful translation for developing efficient drug against cancer cells.

Guanine-rich oligonucleotides (GROs) can self-assemble into four stranded G4 structures stabilized by π-π stacking between G-quartets and *via* Hoogsteen hydrogen bonding. These G4 quadruplexes are found for instance in telomeric and promoter regions, where they participate in a large diversity of biological processes. For example, the control region of the proto-oncogene *KRAS* contains a nuclease hypersensitive element (NHE), which can bind to nuclear proteins and is known to form G-quadruplex structures. Also, it is known that parallel-stranded G-quadruplex stabilized in the c-*Myc* promoter region functions as a transcriptional repressor element ^19,20,21,22^. Likewise, the *c-KIT* ^23,24^ and *BCL2* ^25^ oncogenes feature parallel G quadruplexes in their promoter regions and could be targeted by different approaches ^26,27,28^.

Previous works have shown that the expression of KRAS in pancreatic cancer cells can be downregulated by using decoy oligonucleotides mimicking one of the potential NHE quadruplexes ^29,30,31,32^. In normal cells, *KRAS* transcription is blocked by the two G-quadruplexes, then it is activated when MAZ (a G4-binding protein) unfolds the G quadruplex structures, thus inducing the formation of transcription complexes (Figure 1A). Thanks to a decoy strategy it was demonstrated that synthetic G4 oligonucleotides (G4-Oligo) inhibit oncogenic KRAS in human cancer cells. However, this promising, “decoy approach”, which could be potentially extended to many G4 proteins involved in quadruplex binding and resolving (*i.e*. telomere and promoter regions, RNA quadruplexes or quadruplexes resolving helicases… etc.) is facing major issues. Hence clinical applications of this decoy strategy are limited by: i) the cellular internalization of the decoy G4-Oligos (additional cationic transfection reagents are required, for example see ref ^29^), ii) the chemical stability of G4-Oligos, and the iii) supramolecular stability G4-Oligos based quadruplexes in the adequate topology inside the cell. The latter is a major hurdle since the G4 supramolecular structures are required for the final biological activity. Hence, therapeutic approaches allowing the inhibition of both *KRAS* and c-*Myc* oncogenes expression would be highly desirable in the field of cancer.

**Figure 1:**
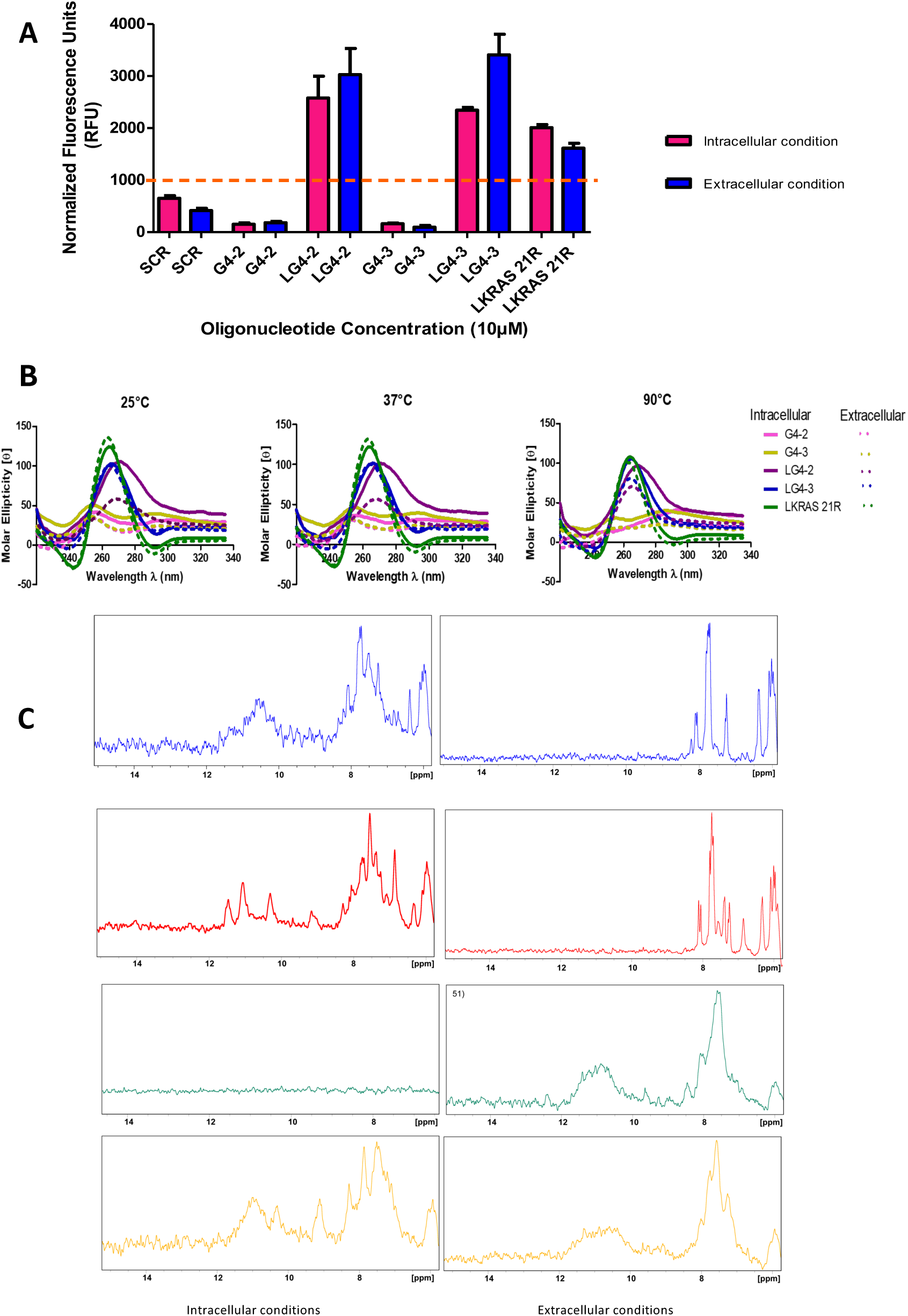
Determination of G quadruplex formation. A- Thioflavin T assay. Enhanced fluorescence of Thioflavin T is observed in the presence of G-quadruplexes under both saline conditions. The threshold for G quadruplex formation is set at 1000 RFUs after normalization using the sequence length. B- Ellipticities of unconjugated and lipid-modified G4 sequences under intracellular and extracellular conditions as determined by circular dichroism. Lipid conjugated G quadruplexes have a negative band at 240 nm and a high ellipticity around 265 nm, characteristic of parallel G quadruplex structure. The topology is more stable under intracellular conditions due to the high concentration of K^+^ ions. C- Imino proton spectrum of the unmodified and NLG4s at physiological saline conditions. G4 formation is characterized by a peak at 10.5 to 12 ppm.

Herein, we propose to explore a "decoy" mode of action by taking advantage of the properties of nucleolipid-oligonucleotides G4 (NLG4-Oligos) (Scheme 1). We hypothesized that the NLG4-Oligos can be active molecules simultaneously on proto-oncogenic targets such as KRAS, MYC, KIT and BCL2. If direct inhibitors targeting each of these oncoproteins are currently being investigated, new strategies, such as the one proposed in this contribution, are needed to simultaneously address these oncogenic targets. Recently, we demonstrated that nucleolipidic tetramolecular parallel G4-Oligos can be formed and manipulated to control both their folding and stability ^33,34^. As a result, it is now possible to obtain tetramolecular parallel G4-Oligos of different length with conformational control and tunable stability. Importantly, we also demonstrated that the lipid modifications were able to improve both the cellular uptake and the *in vivo* delivery of oligonucleotides ^35^.

Building on our previous findings, we postulated that this next-generation class of tetramolecular parallel G-quadruplexes could structurally emulate the endogenous quadruplex motifs within the *KRAS*, *MYC*, *KIT* and *BCL2* oncogene promoters, thereby acting as G4 decoys capable of concurrently and simultaneously suppressing their transcriptional activity. The introduction of lipid moieties facilitates intracellular delivery through the formation of highly stable micellar assemblies, maintaining the integrity of the parallel topology. Collectively, these findings establish a robust proof of concept that NLG4-Oligos can be effectively leveraged as a versatile platform for G4-decoy–based therapeutic strategies.

## 2. RESULTS

### Lipid conjugation of the G4 oligonucleotides enables their self-assembly in micellar structures

In this study, we hypothesized that synthetic stable parallel G4 could act as intracellular decoys of natural G4 sequences. Thus, the oligonucleotides sequences used were chosen according to literature (KRAS21R) ^36^, Hotoda’s sequence and in house developed sequences were synthetized with PTO backbone (Table S1, Figures S1 and S2). To facilitate their internalization in cells, oligonucleotides were conjugated at the 5′-end with a Ketal NucleoLipid phosphoramidite (KNL). For each oligonucleotide, KNL modified and non-modified sequences were compared. As we previously demonstrated, lipid modified oligonucleotides self-assemble in micelles and larger objects ^37,34,33^. The mean size of micellar population of the different G4 sequences was measured by Dynamic Light Scattering (DLS). In both extracellular and intracellular conditions, ranged around 13-26 nm in diameter, with negative zeta potential as expected regarding polyanion structure of oligonucleotides (Figure S3). When the oligonucleotides are resolved in agarose gel, the LONs with guanine-rich sequences form clear bands with a higher molecular weight than their non-lipidic counterparts (Figure S3). There is no significant difference between the bands of the short LONs under high concentrations of potassium (intracellular condition) and sodium ions (extracellular condition). However, a clear difference in molecular weight is observed with the Lipid KRAS 21R, with a retarded band at high potassium ion concentration as opposed to the one under high levels of sodium ions.

**Scheme 1.**
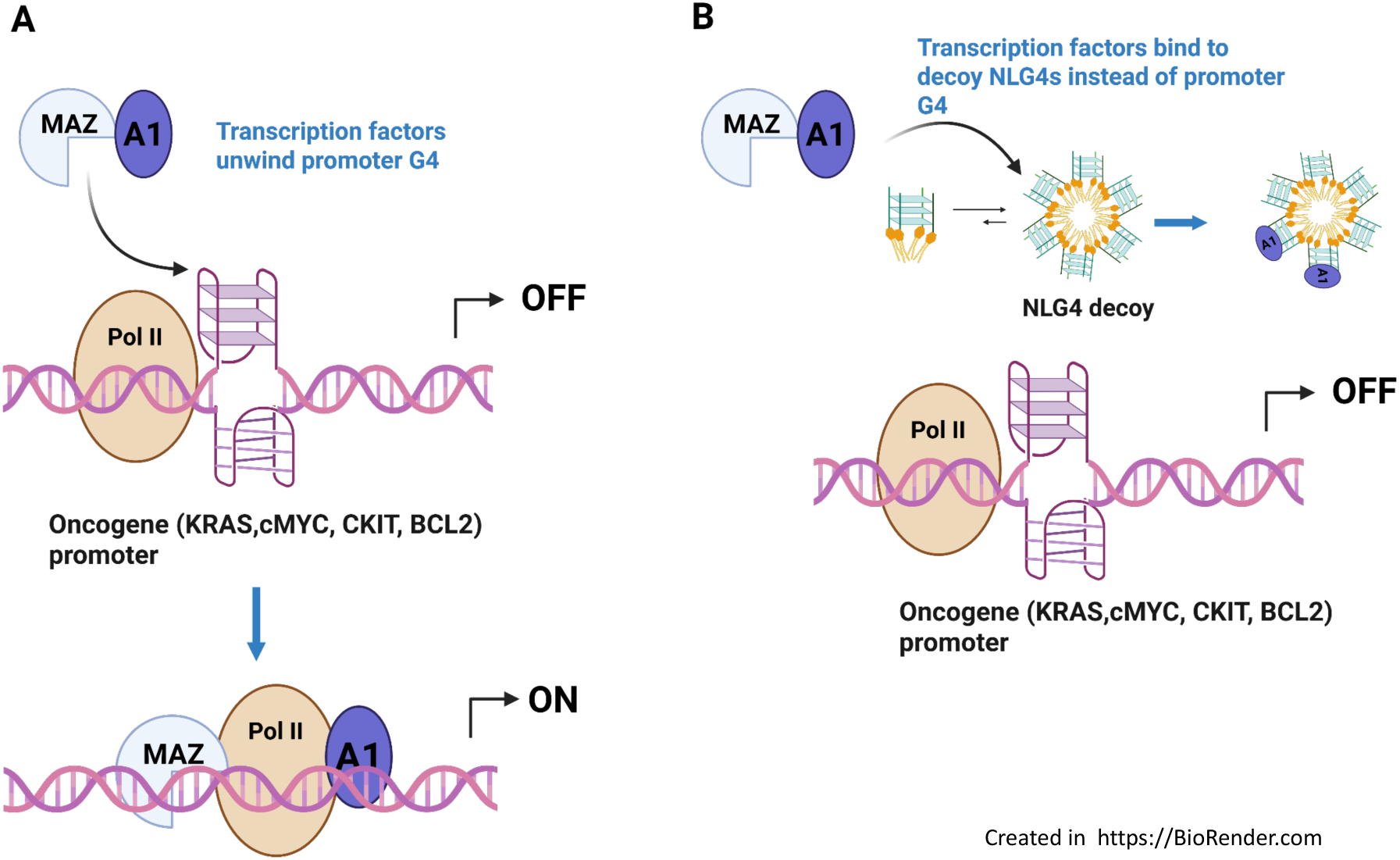
The G4 *decoy strategy*. A- In normal conditions, *KRAS, MYC, KIT and BCL2* transcription are blocked by the G-quadruplexes (OFF). Transcription is activated when MAZ (and cofactor A1) binds to and unfolds the G-quadruplexes (*KRAS*, *c-MYC, KIT and BCL2,* ON), whereas in the presence of decoy G4 MAZ binds to the decoy G4 preventing the transcription of the oncogenes (OFF). B- In this work, we propose to take advantage of the physicochemical properties of the NLG4 (Lipid conjugated parallel tetramolecular G4). NLG4 deliver inside the cell the stable parallel G4, allowing a strong binding to the zinc-finger transcription factors (MAZ and A1) necessary to activate *KRAS, MYC, KIT and BCL2* expression. As a result, in the presence of NLG4 decoys the transcription of the oncogenes are maintained in a off state.

### Conjugation of lipids to guanine-rich sequences facilitates the formation of G quadruplexes that have a tetramolecular parallel topology and are thermal stable

Thioflavin T assay was used to determine the formation of G-quadruplexes (G4s) in both KNL oligonucleotide conjugates and unconjugated guanine-rich sequences. Thioflavin T, a cationic benzothiazole, binds specifically to G-quadruplexes resulting in enhanced fluorescence, while a lower fluorescence increase is observed for single and double-stranded sequences. The fluorescence of the short unconjugated oligonucleotides was lower in contrast to the self-complementary non-G-quadruplex forming sequence used as a control. With the threshold of fluorescence intensity set at 1000, short guanine-rich LONs and the Lipid KRAS 21R had a higher fluorescence at both intracellular and extracellular conditions confirming the formation of G4s (Figures 1A and S4A). The short LONs formed tetra-molecular G4s that resulted in a more than 200-fold increase in fluorescence intensity, compared to the unconjugated sequences. Circular dichroism, CD, was carried out to confirm the formation and topology of G4s. The non-lipidic oligonucleotides had a positive band at 255 nm regardless of the saline conditions, with their lipid-conjugated counterparts having a negative band at 240nm and a positive one at 260nm, a characteristic of parallel G4s (Figure 1B). Hence, lipid conjugation resulted in the formation of G-quadruplexes, in a parallel conformation. The Lipid KRAS 21R sequence had a similar peak profile to the short guanine-rich LONs. CD melting experiments show that lipid conjugation enhances the thermal stability of the G4s formed under both intracellular and extracellular conditions as all LONs maintain their peak profile under the different temperatures (Figure S4B). A NMR study of the G4 formation, which is characterized by several imino peaks in the 10.5 to 12 ppm region, was observed with the unmodified and KNL modified G4s at physiological saline conditions. Despite strong spectral-widths for the imino peaks, all samples show the formation of G4 fingerprint. All together the data collected allow us to propose a G4 architectures reassembled in figure 2.

**Figure 2:**
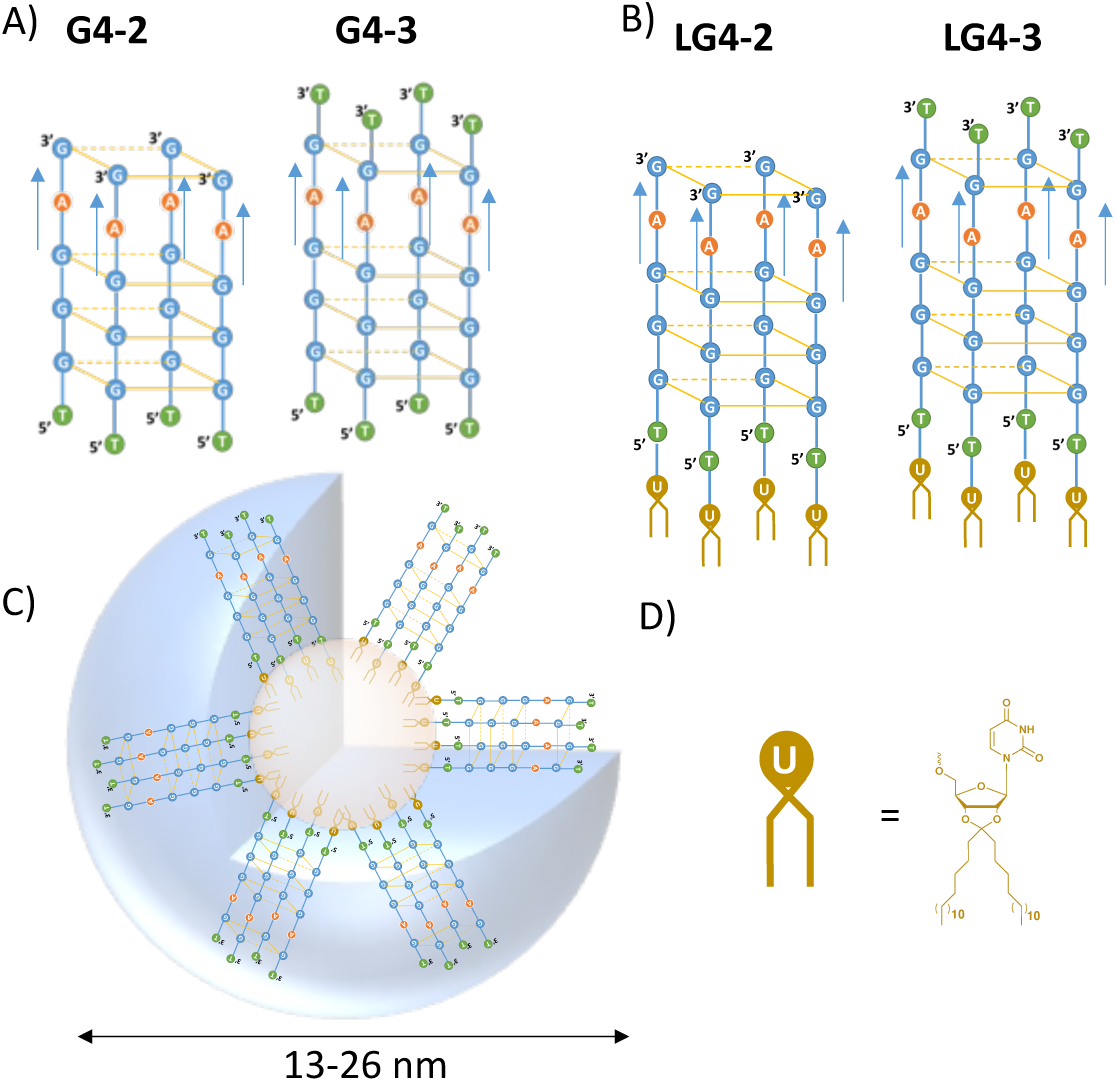
Overview of the G4 structures used in this study. A- Drawing showing a tetramolecular parallel G4 oligonucleotides formed by the sequences G4-2 and G4-3 formed in intracellular conditions. B- Ketal Nucleolipid conjugates (LG4-2 and LG4-3) stabilizing the tetramolecular parallel G4 architecture observed in both extra and intracellular conditions. C- Illustration of the micellar assemblies featuring a hydrophobic core (brown) surrounded by the parallel G4. These assemblies result from the self-aggregation of the tetramolecular lipid parallel G4 (LG4-2 and LG4-3) in both extra and intracellular conditions. D- Chemical structure of the Ketal Nucleolipid (KNL) moiety inserted at the 5’ extremities via a 5’-5’ PS linkage.

In order to determine the possible interactions of the G4 structures electrophoretic mobility shift assays were performed in either the presence or absence of the UP1 protein, a transcription cofactor^38^. In this experiment the oligonucleotides are incubated in the absence (-) and presence (+) of GST tagged UP1. As shown in figure S4-C, the Lipid G-quadruplexes interact with the UP1 protein hence retarded on the gel.

### NLRG4 treatments decrease KRAS, cMYC, and cKit key oncogene levels in pancreatic cancer cells

We next evaluated the efficacy of our G4 decoy oligonucleotides to inhibit the transcription of the oncogenes displaying G4’s structures in their promoter regions like KRAS, cMYC and cKit. We found that in the HPAFII and AsPC-1 PDAC cell lines, the G4s oligonucleotides enable the reduction in key oncogenes protein production when conjugated with lipids (LG4.2 and LG4.3; Figure 3). In HPAFII cells, we observed a 60% decrease in KRAS, cMYC and cKIT protein levels after 72h of LG4.2 treatment (Figure 3 A and B). LG4-3 treatments induced a 20% reduction in KRAS and cKIT production and no significant decrease in cMYC protein levels in HPAFII cell line (Figure 3 A and B). In AsPC1 PDAC cells, LG4-2 treatment induced a 50%, 70% and 20% decrease in KRAS, cMYC and cKit protein levels respectively (Figure 3C and D). In this cell line, LG4.3 oligonucleotide displayed a better efficacy compared to the HPAFII cells with a reduction of 40%, 70% and 25% in the key oncogenes production (Figure 3 C and D).

**Figure 3:**
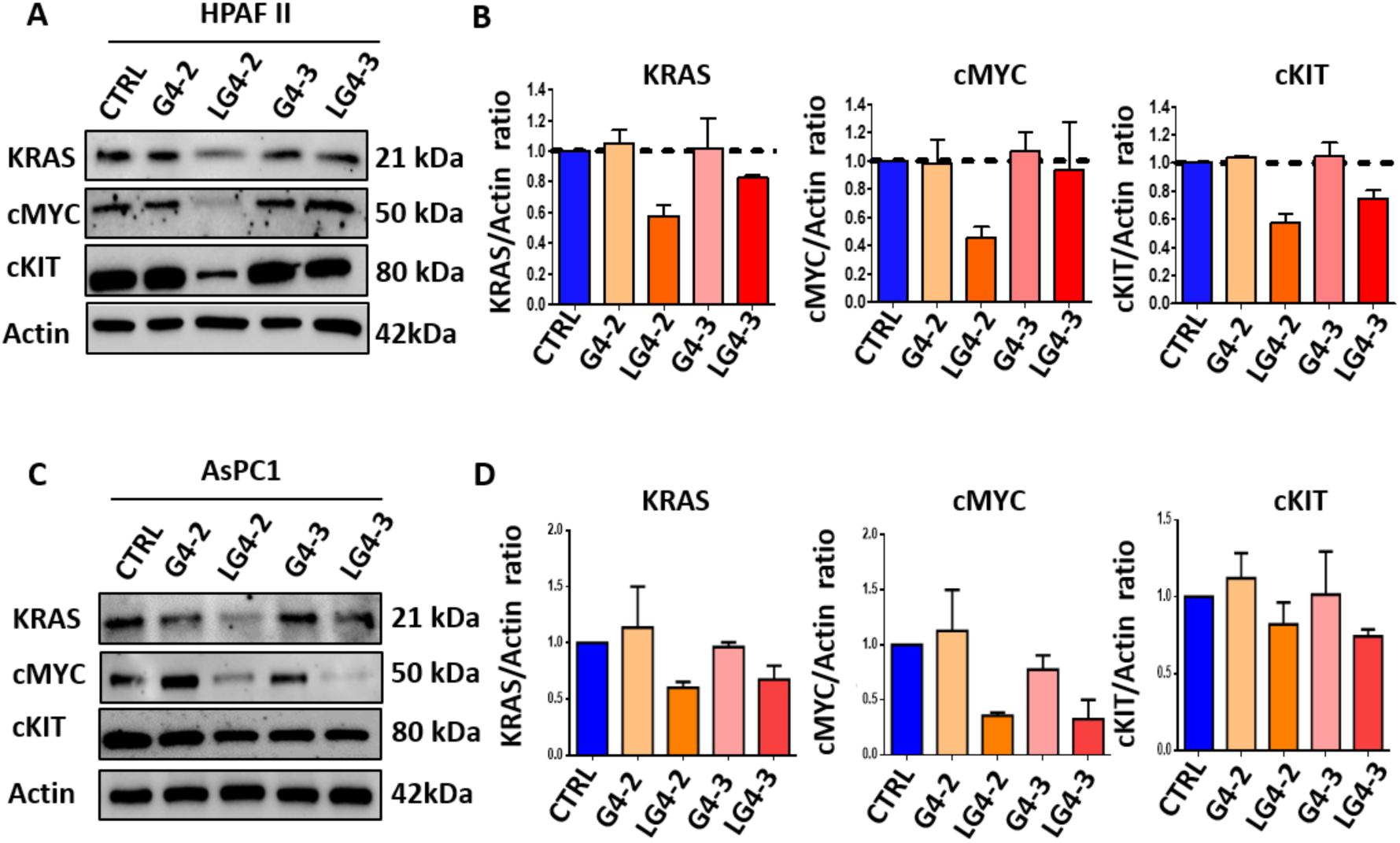
G4s decoys treatments prevent KRAS, cMYC and cKIT oncogenes expression in PDAC cells. A and C- Representative images of western blots showing oncogenes protein levels in HPAFII (A) and AsPC-1 (C) PDAC cell lines after indicated treatments for 72 hours at 10uM (CTRL = Control G4 Oligonucleotide). Actin was used as the house keeping gene. B and D- Quantification of the western blots performed on 3 independent experiments, using ImageJ software.

### Pancreatic cancer cells growth is reduced after lipid-conjugated tetra molecular G quadruplexes treatment

We further questioned the impact of our lipid-G4 oligonucleotides (LG4.2 and LG4.3) on PDAC cell viability and survival markers Ki67 and BCL2 levels. The viability of HPAFII (Figure S5), AsPC1 and BxPC3 PDAC cell lines after G4 decoys treatment for 72h at 10µM. The growth of the three cell lines was significantly reduced in LG4.2 and LG4.3 treated cells (Figure 4 A and C, Figure S6A). LG4.2 treatment induced a 35%, 50% and 45% inhibition of HPAFII, AsPC1 and BxPC3 cells viability respectively. Significant decreased cell viability was also observed in PDAC cells after LG4.3 treatments illustrated by a 60%, 38% and 25% reduction in HPAFII, AsPC1 and BxPC3 cell lines growth respectively (Figure 4 A and C; Figure S6A). Consistent with these observations, LG4.2 and LG4.3 treated PDAC cells expressed low levels of the cell proliferation Ki67 and anti-apoptotic BCL2 markers (Figure 4 B and D). Altogether, these results suggest that our G4 decoy strategy prevent the transcription and production of key oncogenes, well described to have pivotal role in PDAC progression and drug resistance, leading to the decrease in cancer cell viability and proliferation rates. Interestingly, we found no or very small impact of LG4.2 and LG4.3 treatments in BEAS-2B normal cell line growth (Figure S6B).

**Figure 4:**
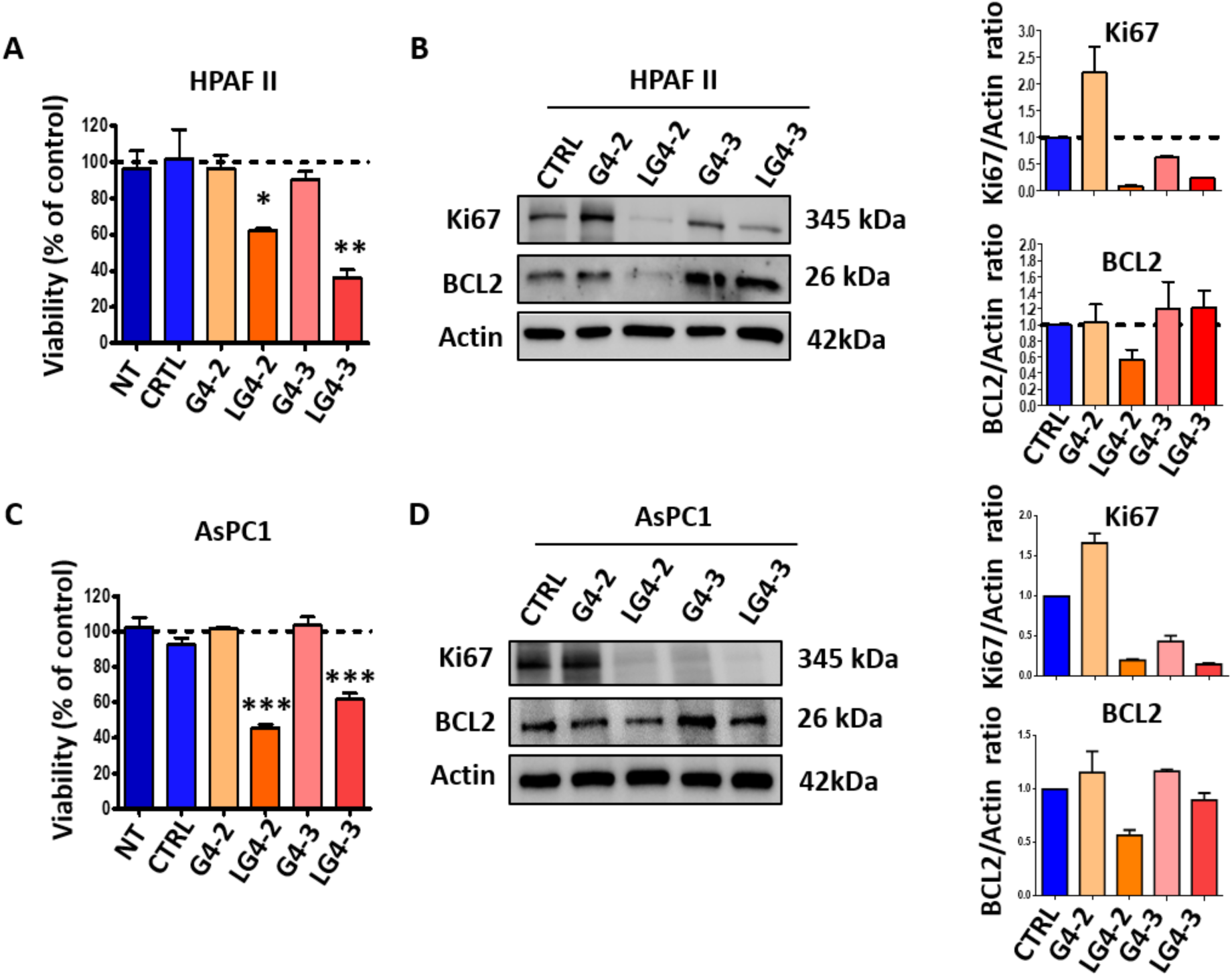
G quadruplex decoy treatments inhibit PDAC cells growth. A and C- Inhibition of HPAF II (A) and AsPC1 (C) cell growth when treated with the lipid-modified G4 oligonucleotides and controls at the concentration of 10µM for 72h. B and D- Representative western blot images reflecting Ki67 and BCL2 protein levels in HPAFII (B) and AsPC1 (D) cells, treated with the indicated conditions at 10 µM. All the experiments were performed in independent triplicate.

### Key markers of canonical oncogenic signalling pathways are reduced in NLG4 treated PDAC cells, leading to the decrease of their migratory and proliferative capacities

NF-kB, Ras/Raf/MEK/ERK and PI3K/Akt signaling are critical for the development and progression of PDAC tumors. The deregulation of these key oncogenic pathways is associated with poor prognosis and drug resistance in PDAC patients. Oncogenes activate these signaling pathways to promote malignant transformation, tumor cell growth, proliferation, migration and apoptosis resistance. We thus interrogated the expression levels of key players involve in oncogenic pathways transduction. In HPAFII cells, LG4-2 treatment reduced by half the levels of NFkB, phospho-Akt (pAkt) and ERK1/2 (Figure 5A). Despite significant reduction in LG4-3 treated HPAFII cell viability, we didn’t observe important changes in oncogenes and signaling key effectors protein levels (Figure 3A and 5A). In AsPC1 cells both NLG4 treatments, LG4-2 and LG4-3, downregulated NFkB, pAkt, Akt and ERK1/2 protein levels (Figure 5D). To further assess the impact of NLG4 treatments on PDAC cells migratory capacities and growth, we performed migration assays and cultured HPAFII, AsPC1 and BxPC3 cell lines in Matrigel™, 72h after the indicated treatments. G4 decoys conjugated with NLs, LG4-2 and LG4-3, significantly reduced HPAF II, AsPC1 and BxPC3 cells migration in comparison to the controls (Figure 5 B and E, Figure S6C). The cell growth abilities of PDAC cells, treated with NLG4 and control oligonucleotides, were interrogated measuring the spheroid diameters 2 weeks after they were embedded in Matrigel™. The average spheroid diameters of lipid conjugated G4 oligonucleotides were significantly decreased in PDAC cells when compared to the non-conjugated G4 oligonucleotide counterparts (Figure 5 C and F; Figure S6D). This last result show that NLG4 based treatments impact on PDAC spheroid growth can last for up to 2 weeks.

**Figure 5:**
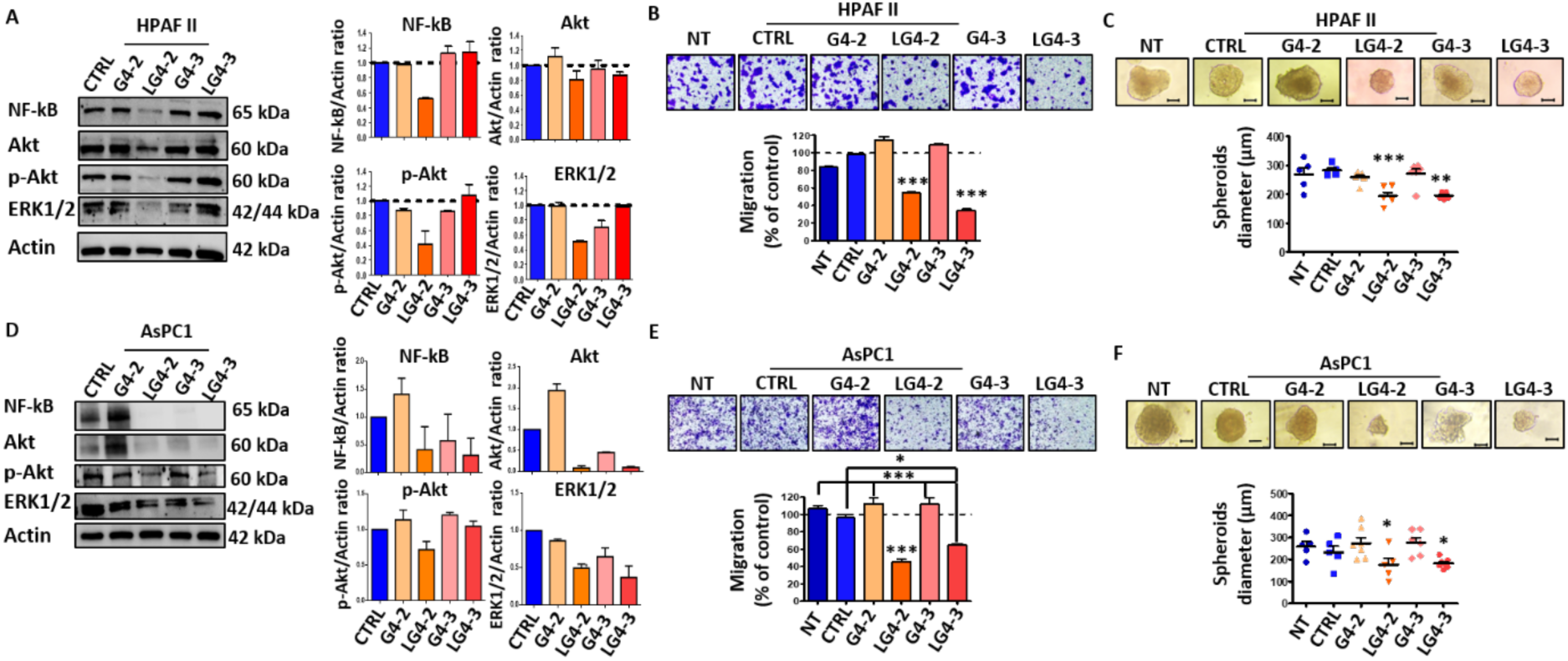
NLG4 treatments reduce critical oncogenic signaling pathway players in PDAC cells, modulating migration and spheroids growth abilities. A and D- Representative western blots showing NF-kB, Akt/p-Akt and ERK1/2 protein levels in HPAFII (A) and AsPC1 (D) cell lines, 72h after the indicated treatments. Western blot quantifications are represented with bar graphs following ImageJ software analyses in three independent experiments. B and E- PDAC cells migration capacities were evaluated using transwells, 72h after the administration of the indicated treatments (NT= Non Treated cells). Representative images of HPAFII (B) and AsPC1 (E) migrated cells stained with crystal violet are showed in the first line (n=3 transwells per treatment). For quantification (graph bars shown in second line), crystal violet was solubilized in ethanol and the absorbance at 555 nm was measured. C and F-Spheroids diameters were measured 2 weeks after HPAFII (C) and AsPC1 (F) cells were embedded in Matrigel™, using the Optika PROView analysis software. Representative images of the spheroids are showed in the first line (scale bar = 100 µm). The quantification was performed on n=6/7 spheroids per conditions. Statistically significant differences between the treatments were determined by analysis of variance (one-way ANOVA). p-values of Tukey’s post-hoc analyses are presented. *P<0.05, **P<0.01 and ***P<0.001.

### NLG4 treatments restore PDAC cells sensitivity to gemcitabine-based chemotherapy

Gemcitabine (GEM) is the standard-of-care chemotherapeutic agent to treat advanced/metastatic pancreatic cancer; however, its clinical benefits are limited due to resistance occurrence. As NLG4-based therapeutics inhibit signaling pathways associated with chemoresistance promotion, we evaluate their ability to restore the PDAC cells sensitivity to gemcitabine. HPAFII, AsPC1 and BxPC-3 cell lines were first treated with G4 oligonucleotides in combination with gemcitabine at 5 µM. The administration of gemcitabine only displayed small effect on G4 controls treated HPAF II cells, with a 15 to 20% decrease in cell viability (Figure 6A). Interestingly, the efficacy of gemcitabine was significantly increased when combined with NLG4-based therapies, leading to additional reduction of 25 and 20% in cell viability following LG4.2 and LG4.3 treatments respectively (Figure 6 A). AsPC1 cells were more sensitive to gemcitabine with a cell viability of 30% in the non-treated (NT) and scrambled oligonucleotide (CTRL) conditions (Figure 6B). However, the combination of G4 decoys with gemcitabine enhanced the chemotherapy cytotoxicity, notably when AsPC1 cells were co-treated with LG4.2 and LG4.3 (Figure 6B). Same results were found in BxPC3 cell line where the combinatorial effect of NLG4-based treatments and gemcitabine led to important reduction of tumor cell viability (Figure S6E). We next investigated the ability of G4 decoys to reduce PDAC cells viability in gemcitabine-resistant cell lines. We induced chemoresistance to HPAFII, AsPC1 and BxPC3 by treating them with increased concentrations of gemcitabine for 3 months. Once the PDAC cells were growing under the chemotherapy pressure, we treated them with G4 oligonucleotides and controls to assess cell viability 72h after the indicated treatments. In HPAFII cells, both LG4.2 and LG4.3-based therapies restored gemcitabine efficacy by reducing significantly cell viability (Figure 6C). In AsPC1 and BxPC3 cell lines, LG4.3 treatments enabled the restoration of gemcitabine sensitivity and LG4.2 had limited impact on cell viability (Figure 6D; Figure S6F).

**Figure 6:**
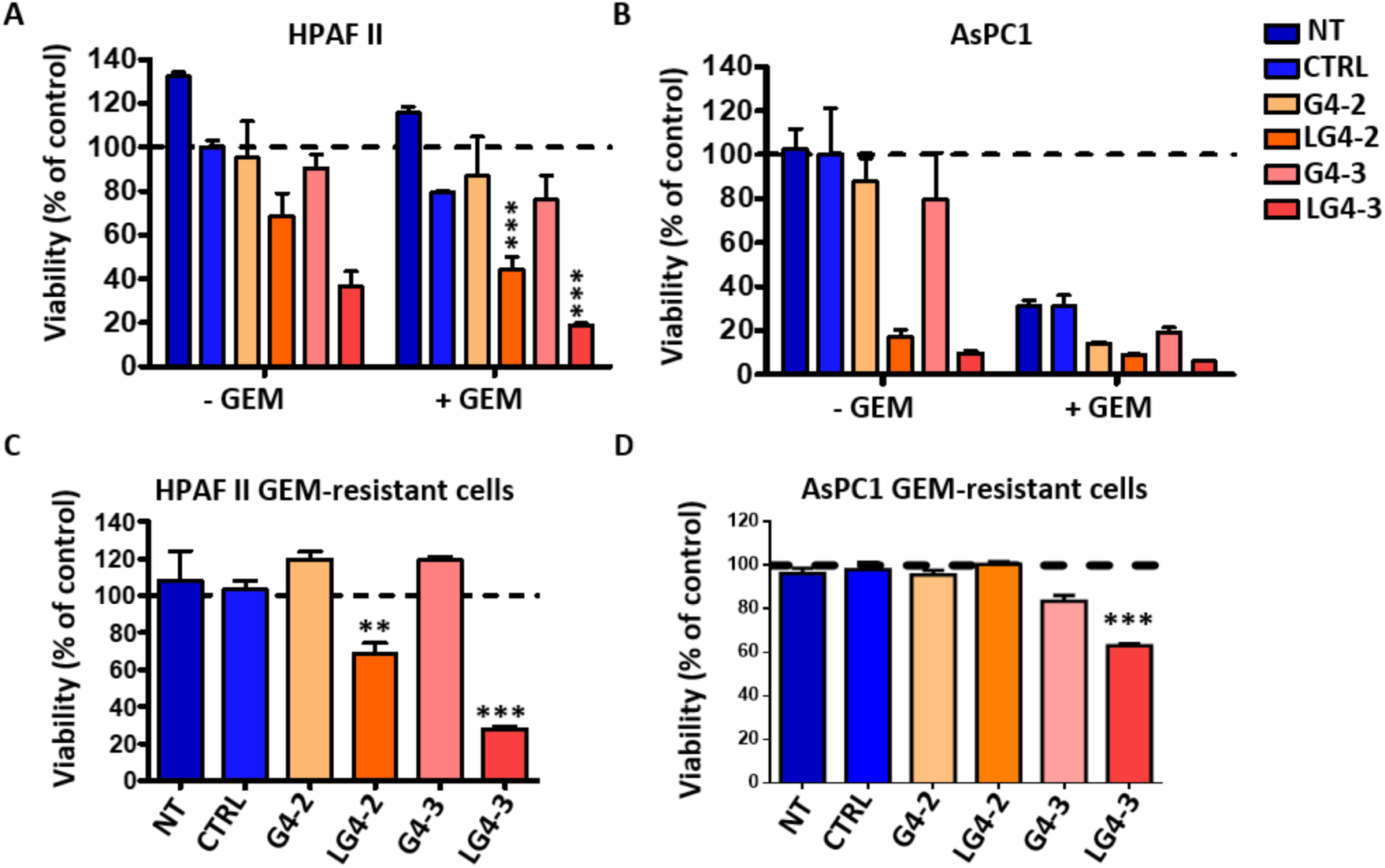
G4 oligonucleotides conjugated with NLs treatments enable the sensitivity restoration of PDAC cells to chemotherapy. A and B- Viability assays were performed on HPAFII (A) and AsPC1 (B) cells treated with G4 oligonucleotides and controls (NT and CTRL) at 10 µM in combination (+GEM) or not (-GEM) with gemcitabine (GEM) chemotherapy at 5 µM for 3 days. C and D- Gemcitabine (GEM) resistant PDAC cell lines were established by intermittent exposure to the chemotherapy. HPAFII (C) and AsPC1 (D) GEM-resistant cell lines were treated with G4 oligonucleotide decoys and control conditions at 10 µM for 72h and the viability of the cells was measured using CellTiter Blue reagent. Statistically significant differences between the treatments were determined by analysis of variance (one-way ANOVA). p-values of Tukey’s post-hoc analyses are presented. **P<0.01 and ***P<0.001.

## 3. DISCUSSION

Several studies indicate that oncogenes, in particular *KRAS* and *c-Myc*, as their mutant alleles are involved in the pathogenesis of different type of human cancers ^39^. KRAS represents the most commonly mutated and MYC the most amplified. Their cooperative interaction constitutes a complex and multifaceted process that profoundly impact tumor initiation and progression. Acting both independently and in concert with other oncogenes, *KRAS* and *c-Myc* orchestrate virtually all the hallmarks of cancer, including immortality, evasion of cell death, tumor-induced angiogenesis, cell invasion and metastasis, metabolic reprogramming, and immune escape. Given their pervasive alterations and central roles in oncogenesis, KRAS, MYC and all their possible related oncogenes or factors stand out as highly compelling targets for cancer therapy.

It is worth emphasizing that most oncogenes are not inherently capable of sustaining uncontrolled cellular proliferation. Rather, they have evolved built-in self-limiting mechanisms that, upon hyperactivation, engage intrinsic cellular fail-safes such as apoptosis or senescence. This principle is exemplified by the prototypical oncogenes KRAS and MYC. Ectopic activation of KRAS, for example, can elicit a cell-cycle arrest phenotype ^40^ indistinguishable from replicative senescence, representing an important barrier that must be bypassed to enable continuous neoplastic growth. In parallel, aberrant MYC expression frequently provokes apoptotic signaling cascades ^41^, necessitating cooperating mutations that suppress programmed cell death to unlock its full tumorigenic potential. In fact, avoiding the oncogenes interplay and tumor progression requires multi-level intervention, by targeting the oncogenes themselves, their cooperative signaling networks, and the tumor ecosystem that sustains them ^42^. Among the promising strategies would be to combine precision medicine ^43^, synthetic lethality ^44^, and immune modulation ^45^ to dismantle the cooperative framework that allows cancer cells to thrive.

Alternatively, one could hypothesize that a concomitant targeting the oncogenes expressions would strongly impact on the cancer cells fate, including tumor growth, cell migration and inflammation of the surrounding tissues. In theory, such an appealing strategy would be possible by targeting concomitantly common DNA motifs localized in promoter regions of theses oncogenes. As a result, this simultaneous targeting would inhibit the transcription steps of these oncogenes and allow to struggle globally against the tumor progression *via* a multiple and efficient oncogenic approach.

**Scheme 2.**
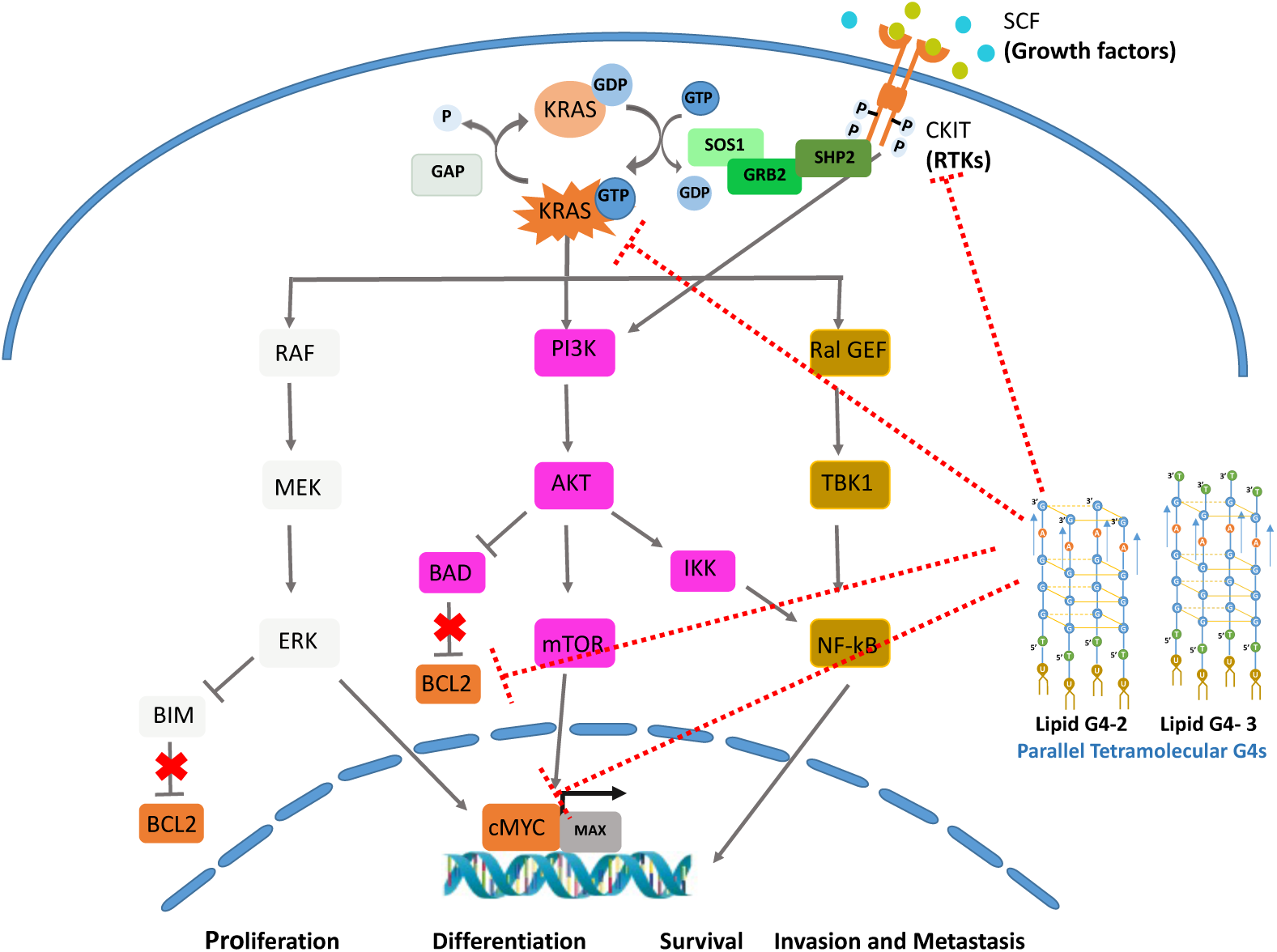
Targeting key oncogenic signaling pathways with NLG4-based treatments. Aberrant activation of KRAS drives PDAC initiation and growth. CKIT is a receptor tyrosine kinase (RTK) that promotes the GDP to GTP switch and activation of KRAS. However, mutations in *KRAS* gene, found in ≈ 95% of PDAC tumors, lead to the constitutive activation of the KRAS protein. Activated KRAS controls downstream pro-oncogenic signaling pathways involved in cancer cells proliferation, survival, differentiation, invasion and metastases. KRAS notably activates *cMYC* transcription *via* the mTOR pathway, and BCL2 release *via* PI3K/Akt and Ras/Raf/MEK/ERK cascade. G-rich sequences were identified in pro-oncogenic promoters including *KRAS*, *cMYC*, *BCL2* and *cKit*, highlighting the importance of G4s in oncogene transcription regulation in cancer cells. To repress the transcription of these oncogenes in PDAC, we developed lipid-modified G4 decoys, LG4.2 and LG4.3, which appears to be an appealing therapeutic approach to prevent drug resistance emergence. In this study, we showed that LG4.2 and LG4.3 enable the downregulation of KRAS, cMYC, BCL2 and cKit simultaneously in PDAC cells, decreasing the level of key effectors of canonical oncogenic pathways and reducing tumor cell survival, growth and migration.

Interestingly, several studies indicate that oncogenes promoters contain G-rich elements capable to fold in G4-DNA. Thus, depending on trans-interacting elements, original G4s present in promoter regions these structures could switch to different topologies giving rise to inhibition or promotion of gene transcription. Hence, the mimicry of such G4 structures using synthetic oligonucleotides would elicit their inhibition *via* specific interaction with natural G4-binding proteins resulting in a decoy mechanism. Thus, synthetic modified oligonucleotide decoys for the concomitant down regulation of these oncogenes would be highly desirable. Among the limitations of such strategies one can note i) the cellular internalization of the decoy structures (G4) and ii) the stabilization of the right G4 topologies in intracellular conditions. As demonstrated by the CD and DLS studies, the lipid modified tetramolecular G4 (Figure 2) allow the intra cellular delivery of decoy oligonucleotides featuring topologies similar to the ones reported for these oncogenes ^20,46,25^. As a result, after entering inside the cancer cells, the decoys elicit their activities on the oncogenes network by a concomitant inhibition of the *CKIT*, *KRAS*, *BCL2*, and *cMYC t*ranscription (Scheme 2). Our investigation on the HPAFII and AsPC-1 PDAC cell lines show the actual inhibitions of those oncogenes (Figures 3 and 4) but also key markers of canonical oncogenic signaling pathways such as NF-kB, Ras/Raf/MEK/ERK and PI3K/Akt, which are critical for the development, migration and progression of PDAC tumors (Figure 5). Importantly, the Ki67 marker, which is an excellent marker to determine the growth fraction of a given cell population is also decreased, indicating that the cancer progression is strongly impacted by the presence of lipid G4.

Pancreatic cancer remains one of the most challenging malignancies to treat, with gemcitabine serving as a standard first-line therapy. However, its clinical benefit is frequently limited due to the rapid emergence of drug resistance^47^. Motivated by this therapeutic challenge, our study focused on several pro-oncogenic promoters including *KRAS, cMYC, BCL2* and *cKit*, identified as the most highly expressed gene in pancreatic cancer. Consistently, the elevated expression of these oncogenes across all pancreatic cancer cell lines might represent targets for improving current drug treatments, including gemcitabine. In this context, cotargeting both the oncogenes’ promoters with their Lipid G4 decoys and gemcitabine becomes appealing to expect synergistic effects in killing the pancreatic cancer cell lines. By killing more than 90% of the cancer cells, the promising results obtained with such combinations on both HPAFII and AsPC1 illustrate the efficacy of this strategy.

## 4. CONCLUSION

Lipidation of the G4-prone oligonucleotide sequences was shown to provide unique properties to G4 structure. First, the lipid oligonucleotide conjugates can stabilize tetramolecular parallel G4 conformers in physiological conditions. Such a structure corresponds to a mimicry of the parallel G4 natural sequences, which are known to control the oncogene transcription /expression of therapeutic relevant targets such as *KRAS, cMYC, BCL2* and *cKit*. In this contribution, we demonstrated the strong potential of the NLG4 strategy against cancer cells, with an impact of the parallel G4 micellar structures on both the expression of oncogenes and malignant cells viability. Notably, treatment with lipid G4-based therapies markedly restored gemcitabine efficacy, as demonstrated by a significant reduction in cancer cell viability. Moreover, LG4 treatment reinstated gemcitabine sensitivity in resistant cancer cells, highlighting the potential of combined therapeutic strategies to counteract tumor resistance and progression. Our approach, which could be adapted to other G4 dependent oncogenes opens promising perspectives in the struggle against cancer.

## DATA AVAILABILITY

The data that support the findings of this study and the source data presented in figures can be found in Supplementary Data.

## ASSOCIATED CONTENT

### Supporting Information

The Supporting Information is available free of charge at https://xxx. Materials and Methods, figures and tables for the results discussed in the paper. Additional experimental results; Mass Spectrometry, HPLC, DLS, Characterization of G4 formation, Cell culture experiments.

## AUTHOR INFORMATION

### Present Addresses

^a^ARNA, INSERM U1212, CNRS 5320, University of Bordeaux, Bordeaux, F-33076, France.

## Author Contributions

P.B conceived the study, designed the experiments with VB and TK, critically interpreted data and wrote the manuscript. B.V. and P.K performed synthesis, purification of oligonucleotides and characterization. Physicochemical experiments were realized by G.S., Z.O, F.K. Cell viability, western blot analysis were achieved by F.K.,V.B. and J.S.G. All authors analyzed and interpreted data and critically read the manuscript. All authors have read and agreed to the published version of the manuscript. ^#^ The first two authors should be regarded as Joint First Authors.

## Funding Sources

The authors acknowledge financial supports from the European Research Council with the grant H2020 Marie Skłodowska-Curie Innovative Training Network “OLIGOMED” (F.K., under grant agreement no. 956070). This work was supported by Inserm Transfert (Optoligo program). This work was supported by la “Ligue Contre le Cancer Aquitaine et Charentes” and partially by the ANR grant for the project SurfMemo ANR-21-CE01-0029.

## Notes

F.K., V. B., T.K., B.V., P. K. and P.B. hold intellectual property rights on uses of G4. P.B. holds equity in NAXIA Discovery. The other authors declare no competing interests.

## ACKNOWLEDGMENT

The authors acknowledge Julie Laguarde and Aurore Guedin-Beaurepaire for their contribution to initial and preliminary cell culture assays.

## ABBREVIATIONS

AKT: Protein Kinase B
BCL2: B-Cell Lymphoma 2
C-MYC: Cellular Myelocytomotosis oncogene
DNA: Deoxyribonucleic Acid
ERK: Extracellular signal-regulated kinases
GDP: Guanosine Diphosphate
GEM: Gemcitabine
GTP: Guanosine Triphosphate
GEF: Guanine Nucleotide Exchange Factor
KIT: proto-oncogene, receptor tyrosine kinase
KRAS: Kirsten Rat Sarcoma virus
MAZ: MYC-Associated Zinc finger
NHE: Nuclease Hypersensitive Elements
TOR: Target Of Rapamycin
PDAC: Pancreatic Ductal Adenocarcinoma
PS: Phosphorothioate
RAF: rapidly accelerated fibrosarcoma
RNA: Ribonucleic Acid
RTKs: Receptor Tyrosine Kinases
UP1: Unwinding Protein 1.

## REFERENCES

(1) Huang, L.; Guo, Z.; Wang, F.; Fu, L. KRAS Mutation: From Undruggable to Druggable in Cancer. Signal Transduct. Target. Ther. 2021, 6 (1), 386. 10.1038/s41392-021-00780-4.

(2) Chen, H.; Smaill, J. B.; Liu, T.; Ding, K.; Lu, X. Small-Molecule Inhibitors Directly Targeting KRAS as Anticancer Therapeutics. J. Med. Chem. 2020, 63 (23), 14404–14424. 10.1021/acs.jmedchem.0c01312.

(3) Zheng, B.-X.; Chen, Z.-X.; Wang, Y.-K.; Dong, J.-P.; She, M.-T.; Zheng, Y.-Y.; Zeng, Y.-X.; Zheng, W.-D.; Long, W.; Wong, W.-L. Translational Suppression of KRAS and NRAS via RNA G-Quadruplex-Targeting Small Molecules for Colorectal Cancer Therapy. J. Med. Chem. 2026, acs.jmedchem.5c03088. 10.1021/acs.jmedchem.5c03088.

(4) Allenson, K.; Castillo, J.; San Lucas, F. A.; Scelo, G.; Kim, D. U.; Bernard, V.; Davis, G.; Kumar, T.; Katz, M.; Overman, M. J.; Foretova, L.; Fabianova, E.; Holcatova, I.; Janout, V.; Meric-Bernstam, F.; Gascoyne, P.; Wistuba, I.; Varadhachary, G.; Brennan, P.; Hanash, S.; Li, D.; Maitra, A.; Alvarez, H. High Prevalence of mutantKRAS in Circulating Exosome-Derived DNA from Early-Stage Pancreatic Cancer Patients. Ann. Oncol. 2017, 28 (4), 741–747. 10.1093/annonc/mdx004.

(5) Schleger, C.; Verbeke, C.; Hildenbrand, R.; Zentgraf, H.; Bleyl, U. C-MYC Activation in Primary and Metastatic Ductal Adenocarcinoma of the Pancreas: Incidence, Mechanisms, and Clinical Significance. Mod. Pathol. 2002, 15 (4), 462–469. 10.1038/modpathol.3880547.

(6) Chen, H.; Song, J.; Wang, Y.; Peng, Z.; Zeng, H.; Zhou, H.; Shang, W.; Chen, Y. Alternative Strategies of Reducing MYC Expression for the Treatment of Cancers. J. Med. Chem. 2026, 69 (2), 744–764. 10.1021/acs.jmedchem.5c02684.

(7) Sodir, N. M.; Kortlever, R. M.; Barthet, V. J. A.; Campos, T.; Pellegrinet, L.; Kupczak, S.; Anastasiou, P.; Swigart, L. B.; Soucek, L.; Arends, M. J.; Littlewood, T. D.; Evan, G. I. MYC Instructs and Maintains Pancreatic Adenocarcinoma Phenotype. Cancer Discov. 2020, 10 (4), 588–607. 10.1158/2159-8290.CD-19-0435.

(8) Parasido, E.; Avetian, G. S.; Naeem, A.; Graham, G.; Pishvaian, M.; Glasgow, E.; Mudambi, S.; Lee, Y.; Ihemelandu, C.; Choudhry, M.; Peran, I.; Banerjee, P. P.; Avantaggiati, M. L.; Bryant, K.; Baldelli, E.; Pierobon, M.; Liotta, L.; Petricoin, E.; Fricke, S. T.; Sebastian, A.; Cozzitorto, J.; Loots, G. G.; Kumar, D.; Byers, S.; Londin, E.; DiFeo, A.; Narla, G.; Winter, J.; Brody, J. R.; Rodriguez, O.; Albanese, C. The Sustained Induction of C-MYC Drives Nab-Paclitaxel Resistance in Primary Pancreatic Ductal Carcinoma Cells. Mol. Cancer Res. 2019, 17 (9), 1815–1827. 10.1158/1541-7786.MCR-19-0191.

(9) Casacuberta-Serra, S.; González-Larreategui, Í.; Capitán-Leo, D.; Soucek, L. MYC and KRAS Cooperation: From Historical Challenges to Therapeutic Opportunities in Cancer. Signal Transduct. Target. Ther. 2024, 9 (1), 205. 10.1038/s41392-024-01907-z.

(10) Jänne, P. A.; Riely, G. J.; Gadgeel, S. M.; Heist, R. S.; Ou, S.-H. I.; Pacheco, J. M.; Johnson, M. L.; Sabari, J. K.; Leventakos, K.; Yau, E.; Bazhenova, L.; Negrao, M. V.; Pennell, N. A.; Zhang, J.; Anderes, K.; Der-Torossian, H.; Kheoh, T.; Velastegui, K.; Yan, X.; Christensen, J. G.; Chao, R. C.; Spira, A. I. Adagrasib in Non–Small-Cell Lung Cancer Harboring a *KRAS^G12C^* Mutation. N. Engl. J. Med. 2022, 387 (2), 120–131. 10.1056/NEJMoa2204619.

(11) Johnson, M. L.; Ou, S. H. I.; Barve, M.; Rybkin, I. I.; Papadopoulos, K. P.; Leal, T. A.; Velastegui, K.; Christensen, J. G.; Kheoh, T.; Chao, R. C.; Weiss, J. KRYSTAL-1: Activity and Safety of Adagrasib (MRTX849) in Patients with Colorectal Cancer (CRC) and Other Solid Tumors Harboring a KRAS G12C Mutation. Eur. J. Cancer 2020, 138, S2. 10.1016/S0959-8049(20)31077-7.

(12) Zhou, C.; Li, W.; Song, Z.; Zhang, Y.; Zhang, Y.; Huang, D.; Yang, Z.; Zhou, M.; Mao, R.; Huang, C.; Li, X.; Wang, J. LBA33 A First-in-Human Phase I Study of a Novel KRAS G12D Inhibitor HRS-4642 in Patients with Advanced Solid Tumors Harboring KRAS G12D Mutation. Ann. Oncol. 2023, 34, S1273. 10.1016/j.annonc.2023.10.025.

(13) Bejar, R.; Zhang, H.; Rastgoo, N.; Benbatoul, K.; Jin, Y.; Thayer, M.; Sheng, S.; Chow, G. K.; Montalvo-Lugo, V.; Marango, J.; Howell, S. B.; Rice, W. G. A Phase 1a/b Dose Escalation Study of the MYC Repressor Apto-253 in Patients with Relapsed or Refractory AML or High-Risk MDS. Blood 2020, 136 (Supplement 1), 45–46. 10.1182/blood-2020-141409.

(14) Xu, Y.; Yu, Q.; Wang, P.; Wu, Z.; Zhang, L.; Wu, S.; Li, M.; Wu, B.; Li, H.; Zhuang, H.; Zhang, X.; Huang, Y.; Gan, X.; Xu, R. A Selective Small-Molecule c-Myc Degrader Potently Regresses Lethal c-Myc Overexpressing Tumors. Adv. Sci. 2022, 9 (8), 2104344. 10.1002/advs.202104344.

(15) Skoulidis, F.; Li, B. T.; Dy, G. K.; Price, T. J.; Falchook, G. S.; Wolf, J.; Italiano, A.; Schuler, M.; Borghaei, H.; Barlesi, F.; Kato, T.; Curioni-Fontecedro, A.; Sacher, A.; Spira, A.; Ramalingam, S. S.; Takahashi, T.; Besse, B.; Anderson, A.; Ang, A.; Tran, Q.; Mather, O.; Henary, H.; Ngarmchamnanrith, G.; Friberg, G.; Velcheti, V.; Govindan, R. Sotorasib for Lung Cancers with *KRAS* p.G12C Mutation. N. Engl. J. Med. 2021, *384* (25), 2371–2381. 10.1056/NEJMoa2103695.

(16) Zhou, Y.; Zeng, C.; Sun, X.; Zhang, J.; Qu, H.; Zhang, X.; Zhou, Y.; Liu, Z.; Wu, X.; Wu, X.; Jiao, X.; Shen, L.; Zhou, Y.; Wang, Y.; Li, J. Activity of Anlotinib in the Second-Line Therapy of Metastatic Gastrointestinal Stromal Tumors: A Prospective, Multicenter, In Vitro Study. The Oncologist 2023, 28 (4), e191–e197. 10.1093/oncolo/oyac271.

(17) Alchoueiry, M.; Mansour, H.; Khabibullin, D.; Han, T.; Chaturantabut, S.; Bzeih, W.; Tang, Y.; Williams, J. F.; Hirsch, M. S.; Priolo, C.; Sellers, W. R.; Henske, E. P. Targeting KIT With Antibody-Drug Conjugates in Chromophobe Renal Cell Carcinoma. Clin. Genitourin. Cancer 2025, 23 (4), 102359. 10.1016/j.clgc.2025.102359.

(18) Croce, C. M.; Vaux, D.; Strasser, A.; Opferman, J. T.; Czabotar, P. E.; Fesik, S. W. The BCL-2 Protein Family: From Discovery to Drug Development. Cell Death Differ. 2025, 32 (8), 1369–1381. 10.1038/s41418-025-01481-z.

(19) Seenisamy, J.; Rezler, E. M.; Powell, T. J.; Tye, D.; Gokhale, V.; Joshi, C. S.; Siddiqui-Jain, A.; Hurley, L. H. The Dynamic Character of the G-Quadruplex Element in the c-MYC Promoter and Modification by TMPyP4. J. Am. Chem. Soc. 2004, 126 (28), 8702–8709. 10.1021/ja040022b.

(20) Siddiqui-Jain, A.; Grand, C. L.; Bearss, D. J.; Hurley, L. H. Direct Evidence for a G-Quadruplex in a Promoter Region and Its Targeting with a Small Molecule to Repress c-*MYC* Transcription. Proc. Natl. Acad. Sci. 2002, 99 (18), 11593–11598. 10.1073/pnas.182256799.

(21) Mathad, R. I.; Hatzakis, E.; Dai, J.; Yang, D. C-MYC Promoter G-Quadruplex Formed at the 5′-End of NHE III 1 Element: Insights into Biological Relevance and Parallel-Stranded G-Quadruplex Stability. Nucleic Acids Res. 2011, 39 (20), 9023–9033. 10.1093/nar/gkr612.

(22) Umek, T.; Sollander, K.; Bergquist, H.; Wengel, J.; Lundin, K. E.; Smith, C. I. E.; Zain, R. Oligonucleotide Binding to Non-B-DNA in MYC. Molecules 2019, 24 (5), 1000. 10.3390/molecules24051000.

(23) Wei, D.; Parkinson, G. N.; Reszka, A. P.; Neidle, S. Crystal Structure of a C-Kit Promoter Quadruplex Reveals the Structural Role of Metal Ions and Water Molecules in Maintaining Loop Conformation. Nucleic Acids Res. 2012, 40 (10), 4691–4700. 10.1093/nar/gks023.

(24) Peterková, K.; Durník, I.; Marek, R.; Plavec, J.; Podbevšek, P. C-Kit2 G-Quadruplex Stabilized via a Covalent Probe: Exploring G-Quartet Asymmetry. Nucleic Acids Res. 2021, 49 (15), 8947–8960. 10.1093/nar/gkab659.

(25) Nambiar, M.; Goldsmith, G.; Moorthy, B. T.; Lieber, M. R.; Joshi, M. V.; Choudhary, B.; Hosur, R. V.; Raghavan, S. C. Formation of a G-Quadruplex at the BCL2 Major Breakpoint Region of the t(14;18) Translocation in Follicular Lymphoma. Nucleic Acids Res. 2011, 39 (3), 936–948. 10.1093/nar/gkq824.

(26) Amato, J.; Pagano, A.; Capasso, D.; Di Gaetano, S.; Giustiniano, M.; Novellino, E.; Randazzo, A.; Pagano, B. Targeting the *BCL2* Gene Promoter G-Quadruplex with a New Class of Furopyridazinone-Based Molecules. ChemMedChem 2018, 13 (5), 406–410. 10.1002/cmdc.201700749.

(27) Teng, F.-Y.; Jiang, Z.-Z.; Guo, M.; Tan, X.-Z.; Chen, F.; Xi, X.-G.; Xu, Y. G-Quadruplex DNA: A Novel Target for Drug Design. Cell. Mol. Life Sci. 2021, 78 (19–20), 6557–6583. 10.1007/s00018-021-03921-8.

(28) Kulshreshtha, N. N.; Barthélémy, P. Lock and Key: Locked G Quadruplexes Could Be the Key to New Modalities in Nucleic Acid Therapeutics. RSC Med. Chem. 2025, 16 (8), 3336–3343. 10.1039/D5MD00142K.

(29) Cogoi, S.; Zorzet, S.; Rapozzi, V.; Géci, I.; Pedersen, E. B.; Xodo, L. E. MAZ-Binding G4-Decoy with Locked Nucleic Acid and Twisted Intercalating Nucleic Acid Modifications Suppresses KRAS in Pancreatic Cancer Cells and Delays Tumor Growth in Mice. Nucleic Acids Res. 2013, 41 (7), 4049–4064. 10.1093/nar/gkt127.

(30) Cogoi, S.; Shchekotikhin, A. E.; Xodo, L. E. *HRAS* Is Silenced by Two Neighboring G-Quadruplexes and Activated by MAZ, a Zinc-Finger Transcription Factor with DNA Unfolding Property. Nucleic Acids Res. 2014, 42 (13), 8379–8388. 10.1093/nar/gku574.

(31) Miglietta, G.; Gouda, A. S.; Cogoi, S.; Pedersen, E. B.; Xodo, L. E. Nucleic Acid Targeted Therapy: G4 Oligonucleotides Downregulate *HRAS* in Bladder Cancer Cells through a Decoy Mechanism. ACS Med. Chem. Lett. 2015, 6 (12), 1179–1183. 10.1021/acsmedchemlett.5b00315.

(32) Cogoi, S.; Jakobsen, U.; Pedersen, E. B.; Vogel, S.; Xodo, L. E. Lipid-Modified G4-Decoy Oligonucleotide Anchored to Nanoparticles: Delivery and Bioactivity in Pancreatic Cancer Cells. Sci. Rep. 2016, 6 (1), 38468. 10.1038/srep38468.

(33) Vialet, B.; Bansode, N. D.; Gissot, A.; Barthélémy, P. Controlling the G-Quadruplex Topology: Toward the Formation of a Lipid Thrombin Binding Aptamer Prodrug. Bioconjug. Chem. 2023, 34 (7), 1198–1204. 10.1021/acs.bioconjchem.3c00170.

(34) Vialet, B.; Gissot, A.; Delzor, R.; Barthélémy, P. Controlling G-Quadruplex Formation via Lipid Modification of Oligonucleotide Sequences. Chem Commun 2017, 53 (84), 11560–11563. 10.1039/C7CC05693A.

(35) Karaki, S.; Benizri, S.; Mejías, R.; Baylot, V.; Branger, N.; Nguyen, T.; Vialet, B.; Oumzil, K.; Barthélémy, P.; Rocchi, P. Lipid-Oligonucleotide Conjugates Improve Cellular Uptake and Efficiency of TCTP-Antisense in Castration-Resistant Prostate Cancer. J. Controlled Release 2017, 258, 1–9. 10.1016/j.jconrel.2017.04.042.

(36) Marquevielle, J.; Robert, C.; Lagrabette, O.; Wahid, M.; Bourdoncle, A.; Xodo, L. E.; Mergny, J.-L.; Salgado, G. F. Structure of Two G-Quadruplexes in Equilibrium in the KRAS Promoter. Nucleic Acids Res. 2020, 48 (16), 9336–9345. 10.1093/nar/gkaa387.

(37) Pokholenko, O.; Gissot, A.; Vialet, B.; Bathany, K.; Thiéry, A.; Barthélémy, P. Lipid Oligonucleotide Conjugates as Responsive Nanomaterials for Drug Delivery. J. Mater. Chem. B 2013, 1 (39), 5329. 10.1039/c3tb20357c.

(38) Ferino, A.; Marquevielle, J.; Choudhary, H.; Cinque, G.; Robert, C.; Bourdoncle, A.; Picco, R.; Mergny, J.-L.; Salgado, G. F.; Xodo, L. E. hnRNPA1/UP1 Unfolds *KRAS* G-Quadruplexes and Feeds a Regulatory Axis Controlling Gene Expression. ACS Omega 2021, 6 (49), 34092–34106. 10.1021/acsomega.1c05538.

(39) Casacuberta-Serra, S.; González-Larreategui, Í.; Capitán-Leo, D.; Soucek, L. MYC and KRAS Cooperation: From Historical Challenges to Therapeutic Opportunities in Cancer. Signal Transduct. Target. Ther. 2024, 9 (1), 205. 10.1038/s41392-024-01907-z.

(40) Stubbs, C. K.; Biancucci, M.; Vidimar, V.; Satchell, K. J. F. RAS Specific Protease Induces Irreversible Growth Arrest via P27 in Several KRAS Mutant Colorectal Cancer Cell Lines. Sci. Rep. 2021, 11 (1), 17925. 10.1038/s41598-021-97422-0.

(41) Edwards-Hicks, J.; Su, H.; Mangolini, M.; Yoneten, K. K.; Wills, J.; Rodriguez-Blanco, G.; Young, C.; Cho, K.; Barker, H.; Muir, M.; Guerrieri, A. N.; Li, X.-F.; White, R.; Manasterski, P.; Mandrou, E.; Wills, K.; Chen, J.; Abraham, E.; Sateri, K.; Qian, B.-Z.; Bankhead, P.; Arends, M.; Gammoh, N.; Von Kriegsheim, A.; Patti, G. J.; Sims, A. H.; Acosta, J. C.; Brunton, V.; Kranc, K. R.; Christophorou, M.; Pearce, E. L.; Ringshausen, I.; Finch, A. J. MYC Sensitises Cells to Apoptosis by Driving Energetic Demand. Nat. Commun. 2022, 13 (1), 4674. 10.1038/s41467-022-32368-z.

(42) Petroni, G.; Buqué, A.; Coussens, L. M.; Galluzzi, L. Targeting Oncogene and Non-Oncogene Addiction to Inflame the Tumour Microenvironment. Nat. Rev. Drug Discov. 2022, 21 (6), 440–462. 10.1038/s41573-022-00415-5.

(43) Singh, D.; Dhiman, V. K.; Pandey, M.; Dhiman, V. K.; Sharma, A.; Pandey, H.; Verma, S. K.; Pandey, R. Personalized Medicine: An Alternative for Cancer Treatment. Cancer Treat. Res. Commun. 2024, 42, 100860. 10.1016/j.ctarc.2024.100860.

(44) Ngoi, N. Y. L.; Gallo, D.; Torrado, C.; Nardo, M.; Durocher, D.; Yap, T. A. Synthetic Lethal Strategies for the Development of Cancer Therapeutics. Nat. Rev. Clin. Oncol. 2025, 22 (1), 46–64. 10.1038/s41571-024-00966-z.

(45) Giri, S.; Lamichhane, G.; Pandey, J.; Khadayat, R.; K. C., S.; Devkota, H. P.; Khadka, D. Immune Modulation and Immunotherapy in Solid Tumors: Mechanisms of Resistance and Potential Therapeutic Strategies. Int. J. Mol. Sci. 2025, 26 (7), 2923. 10.3390/ijms26072923.

(46) Burge, S.; Parkinson, G. N.; Hazel, P.; Todd, A. K.; Neidle, S. Quadruplex DNA: Sequence, Topology and Structure. Nucleic Acids Res. 2006, 34 (19), 5402–5415. 10.1093/nar/gkl655.

(47) Quiñonero, F.; Mesas, C.; Doello, K.; Cabeza, L.; Perazzoli, G.; Jimenez-Luna, C.; Rosa Rama, A.; Melguizo, C.; Prados, J. The Challenge of Drug Resistance in Pancreatic Ductal Adenocarcinoma: A Current Overview. Cancer Biol. Med. 2019, 16 (4), 688–699. 10.20892/j.issn.2095-3941.2019.0252.

