## Supplementary material for "Self-Assembled Nucleolipid G-Quadruplexes Act as Multitarget Decoys for Oncogene Suppression in Pancreatic Cancer": Suplementary information

### Materials and Methods

#### Synthesis, purification and dosage of oligonucleotides and Lipid conjugated antisense oligonucleotides

The oligonucleotide synthesis was performed on an automated H8 synthesizer (K&A labs, Germany) at the µmol scale on CPG (controlled pore glass) 1000Å primer support using the standard β-phosphoramidite methodology. All sequences were synthesized with full phosphorothioate linkage and the double-chain nucleolipid (ketal-bis-$C_{15}$-Uridine) was inserted at the 5' end. The sequences are described in Table S1. A sequence that does not form a stabilized G4 structure was used as control (G4 control - CTRL). Lipid-G4 were purified by HPLC, non-lipidic G4 were desalted. All oligonucleotides were characterized by ESI mass spectrometry (Supplementary Table SI1).

#### Oligonucleotides' purification

*Chromatographic analysis of purity.*

All synthesized oligonucleotides were analyzed by using High Performance Liquid Chromatography (HPLC) on Elite LaChrom (VWR) system with a diode array detector at 260nm. Aqueous mobile phase **A** contained 95% of 100 mM triethyl ammonium acetate (TEAA) at pH 7 and 5% of acetonitrile (ACN). Organic mobile phase **B** was composed of 20 % of TEAA 100mM and 80% of ACN. Mobile phases' gradients with increasing the percentage of the organic phase **B** allowed oligonucleotide elution. The analysis runs were 15 minutes.

For non-lipidic oligonucleotides, hydrophobic column Xbridge oligonucleotide BEH $C_{18}$ (Waters) with particles' size of 2.5 µm, 130 Å of porosity and 4.6 x 50 mm of geometry was used. The flow was 2.4ml/min starting with 0% **B** to 30% **B** in 10min, remained for 2 minutes at 30% **B** then returned to initial conditions.

For lipid modified oligonucleotides, Nucleosil $C_4$ column with 4*250 mm geometry and particles size of 5 µm, 300 Å of porosity (Macherey Nagel) was used with a flow of 1.0ml/min. Starting with 0% **B** to 100% **B** in 10min, remained for 2 minutes at 100% **B** then returned to starting conditions.

Purification for lipid oligonucleotides was performed using preparative HPLC method with column XBridge Protein BEH $C_4$ OBD Pre with 30x50 mm of geometry, particles size of 5 µM and porosity of 300 Å. The flow was 56.25 mL/min with 4 minute purification run. Starting from 0% **B** to 100% **B** in 2 min, remained for 1 min at 100% **B**, then went back to initial conditions.

*Desalting methods*

An ultra-filtration system was used to desalt the non-lipidic samples. The columns of Vivaspin Turbo 4 (Sartorius, cut-off 3.5 kDa, membrane Polyethersulfone) were used for oligonucleotides' desalting. Membranes were rinsed with distilled water and then samples were added into the column before being centrifuged at 3000 rpm in 30 min. Three washings were made by adding 2 mL of distilled water into the superior part of the tube and then re-centrifuged as previously. 500 µL of distilled water was added on the membrane to re-suspend oligonucleotide and collect it. Then the membrane was rinsed 3 times with 500 µL of distilled water to limit the waste.

After HPLC purification, the lipidic oligonucleotides were desalted using a regenerated cellulose dialysis membrane (Spectra por 6, cut off-1kD) against distilled water.

#### Oligonucleotides assay

The concentration of all ASOs and LASOs was determined by spectrophotometry Nanodrop® (Thermo Scientific™) at 260 nm with automatic oligonucleotide detection mode.

### Mass analysis

ESI mass spectrometry analyses were carried out on a Thermo Fisher Q-Exactive. Oligonucleotide samples were prepared using 3.5 kD vivacon membrane (Sartorius) for dialysis against 50 mM ammonium acetate (Sigma-Aldrich). Spectra show multi-charged ions (M-z)/z obtained in the negative mode. Measured experimental monoisotopic masses were determined using the value of the first peak of the isotope distribution.

#### Dynamic Light Scattering

The size of particles was determined using the Zetasizer 3000 HAS MALVERN. Samples at a final concentration of 50µM were prepared under extracellular (Phosphate buffer $Na^+$ 200 mM pH 7.2, 1.45 M NaCl) and intracellular (Phosphate buffer $K^+$ 200 mM pH 7.2, 1.4 M KCl,120 mM NaCl, 10mM $MgCl_2$) conditions respectively in 1X buffer and salt concentration, denatured at 90°C for 5min and left at room temperature. Measurements were taken at 25°C.

#### Circular Dichroism (CD)

CD spectra and CD-monitored melting curves were recorded on a JASCO 1500 spectrometer equipped with a Peltier-type temperature control system. CD spectra were recorded from 220 nm to 335 nm at 25 °C with a data pitch of 0.2 nm, a bandwidth of 2 nm, a response of 0.5 s, and a scanning speed of 100 nm/min. The resulting spectra were the averages of three accumulations. The oligonucleotide samples were prepared with 10 mM phosphate buffer at 10 µM at different saline concentrations. The sample solutions were annealed for 5 min at 90 °C, then kept at RT for 2h before analysis.

CD melting experiments were performed overnight from 5 °C to 90 °C. Melting curves were extracted at 265 nm. The holder was used as the control sensor and the cell was used as the monitor sensor. A temperature gradient of 0.5 °C/min and a data pitch of 0.5 nm were used. The scanning speed was 200 nm/min.

#### Thioflavin T Assay

Samples at a final concentration of 10µM were prepared under extracellular (Phosphate buffer Na+ 200 mM pH 7.2, 1.45 M NaCl) and intracellular (Phosphate buffer $K^+$ 200 mM pH 7.2, 1.4 M KCl,120 mM NaCl, 10mM $MgCl_2$) conditions respectively in 1X buffer and salt concentration, denatured at 90°C for 5min and left at room temperature two hours. 20 µL of the sample were mixed with 20µL of 400nM Thioflavin T in a black 96 well plate and fluorescence readings were taken at an excitation and emission wavelength of 440 nm and 490 nm respectively. Fluorescence intensity was normalized by sequence length and analysis was done using GraphPad Prism

**Electrophoretic Mobility Shift Assay**

Oligonucleotide samples at 100µM concentration, were prepared under intracellular saline conditions for G quadruplex formation. A volume of 3µl of the oligonucleotide, 3µl unfolding protein 1 tagged to GST (UP1+ GST tag) at 100µM and 2.6 µl of 5mM 1,4 Dithiothreitol (DTT) were added in a well-labeled Eppendorf tube and incubated at room temperature for 30 minutes. Loading dye at a volume of 2µL was mixed with the contents and resolved on 1% agarose gel in 0.5X Tris Borate (TB) buffer, containing 10µM of Thioflavin T, at 50 V for 1 hour 40 min in the dark. The gel was then imaged using ChemiDoc™ (Bio-Rad)

**Recombinant Protein Production**

GST tagged U1P was recombinantly expressed in E. coli BL21 (DE3) bacteria with LB medium (5 g/L yeast extract, 10 g/L peptone, and 10 g/L NaCl) at 37 °C overnight with ampicillin at 100 µg/mL. Expression was induced at an OD 600 nm between near 0.6-0.8 with 1 mM IPTG overnight at 17 °C. The bacterial pellets were then concentrated by centrifugation at 6500 rpm for 20 min at 4 °C. After which they were resuspension with PBS, and the solution was put under agitation for 30 min with 100 mM PMSF, lysozyme, DNAse, RNAse and 1 M DTT. A lysis by sonication (40%: 45 s on, 45 s off for 4 min and 30 s) was then carried out, the lysate was then ultracentrifuged for 1 h at 4 °C at 42,000 RPM, and the supernatant was collected. Glutathione Sepharose 4B (GE Healthcare) (50% slurry in PBS) was added to the supernatant and incubated for 2 h at 4 °C with a slow shaking. The mix was centrifuged at 2000 rpm for 5 min, and the pellet was washed five times in PBS and eluted with an elution buffer containing 20 mM NaCl, 20 mM reduced glutathione, and 200 mM Tris–HCl (pH 7.5). The purity was checked by 1D NMR and SDS-PAGE and concentration was determined by UV-vis at 280 nm.

**Cell culture**

HPAF II cells (purchased in ATCC, CRL-1997™) were maintained in Minimum Essential Media (Gibco) supplemented with 10% foetal bovine serum (FBS). BxPC3 and AsPC1 cell lines were purchased at the ATCC (CRL-1687™ and CRL-1682™ respectively) and cultured in RPMI (Gibco). BEAS-2B non cancerous cell line was kindly provided by Dr Jeanne Leblond-Chain (ARNA Lab, Bordeaux FR) and cultured in DMEM (Gibco) supplemented with 10% FBS. All the cells were cultured at 37°C and 5% $CO_2$ under humified conditions.

**Cell Viability**

HPAF II, AsPC1 and BEAS-2B cells were seeded in 96-well plates at a density of 5000 cells/well in 100 µL of culture media. BxPC3 cell line was seeded at 2500 cells/well. After 24 hours, the cells were incubated in serum-free MEM, DMEM or RPMI culture media containing oligonucleotides at the final concentrations of 1µM, 5 µM, and 10 µM (transfection media), for 4 hours. Culture media containing 20% FBS was then added to each well, at the volume of 100 µL. Three days later, cell viability was measured using CellTiter® Blue cell viability assay (Promega) following the manufacturer's protocol. Briefly, 40ul of CellTiter® Blue reagent was added to each well and incubated at 37°C for two hours. Fluorescent readings were measured ($\lambda_{exc}$ = 560 nm, $\lambda_{em}$ = 590 nm, 10 nm bandwidth).

**Development of Gemcitabine Resistance Cell Lines**

PDAC cell lines were exposed to increasing concentrations (from 5nM to 5µM) of gemcitabine (Sigma) for 4 months. The gemcitabine concentrations were increased weekly (every 2 passages). Once the 5µM resistant cell lines were established, they were passaged 3 times with the highest concentration of gemcitabine before performing the survival assays.

**Western blot**

Pancreatic tumor cells were plated at the density of 200,000 cells per 10 cm culture plates in 10 mL of culture media for 24 hours. The culture media was replaced with 5 mL of transfection media at a final concentration of 10 µM oligonucleotide for 4 hours. Then, 5 mL of culture media supplemented with 20% FBS was added to each plate. After 3 days, cells were harvested and lysed using Pierce™ RIPA Buffer (Thermo Scientific) following the manufacturer's instructions. Protein quantification was measured using the bicinchoninic acid protein assay kit (Thermo Scientific) and 10 µg of protein per lane were loadedon 15% SDS PAGE gel. After migration, the proteins were transferred to a PVDF membrane. Blocking of the membranes with 5% non-fat dry milk in Tris-buffered saline containing 0.05% Tween 20 (TBST) was done for one hour at room temperature. The membrane was incubated with primary antibodies, for 1-2 hours at room temperature, followed with secondary antibody incubations for 1 hour. SuperSignal™ West Pico PLUS chemiluminescence western blot substrate (Thermo Scientific) was used to reveal the blot. The following antibodies, purchased from Invitrogen, were used: KRAS recombinant rabbit monoclonal antibody (11H35L14, Invitrogen), cMYC recombinant rabbit monoclonal antibody (9E10, Invitrogen), c-Kit recombinant rabbit monoclonal antibody (NH34LC14, Invitrogen), Akt rabbit monoclonal antibody (D6G4, Cell Signalling Technology), Bcl2 recombinant rabbit monoclonal antibody (JF104-8, Invitrogen), Actin recombinant rabbit monoclonal (JF47-01), and goat anti-rabbit poly HRP.

**Migration assays**

Migration assays were performed using Thermo Scientific™ Nunc™ polycarbonate cell culture inserts (membrane pore size of 8 µm) in 24 well plates. After treatments with G4 oligonucleotides at 10 µM, 75 000 cells/well were plated onto the transwell upper chamber in 200 µl of serum free culture media and 500 µl of complete media were added onto the bottom chamber. After 24 hours, the cells were washed twice in PBS, and fixed in PBS + 4% of paraformaldehyde (PFA) for 15 minutes at RT. After 2 washing steps in PBS, the non-migrated cells on the upper surface of the transwells were gently removed with cotton swabs. The migrated cells on the bottom surface of the transwells were stained with 0.2% (w/v) crystal violet for 5 minutes. Migrated cells were observed with a phase-contrast microscope. For the quantification, crystal violet coloration was solubilized in ethanol for 5 minutes, transferred in clean 96 well plates and the optical density was read at 555 nm using a plate reader (Biorad).

**Spheroids culture**
Three days after the transfection with 10uM of oligonucleotides, pancreatic tumor HPAFII, AsPC1 and BxPC3 cells were harvested, pelleted and suspended in Matrigel™ (Corning) at a final concentration of 50 000 cells/ml. The mixture cells/ Matrigel™ was transferred into 96 well plates (100 µl/well) and left at 37°C for 15 minutes to allow the gelation of the matrices. Then, 200 µl of culture media were added on each well and refreshed twice a week. Pictures of the spheroids were taken three weeks after the cells were embedded in Matrigel™, with an inverted microscope (Magnification 10X - Optika). Spheroid diameters were measured two weeks after the seeding, using the Optika PROView analysis software.

**Statistical analysis**
Results were analyzed and represented as mean ± SEM. Statistical analyses were performed using GraphPad Prism 6 software (GraphPad Software, Inc). The one-way ANOVA followed by Tukey-Kramer *post-hoc* test was used to compare the means between the different conditions of culture. Results with $P < 0.05$ were considered statistically significant and are indicated by *, $P < 0.05$; **, $P < 0.01$; and ***, $P < 0.001$.

**NMR spectroscopy**
NMR experiments were recorded on a Bruker Advance III 700 spectrometer equipped with a liquid TXI 1H/13C/15N/2H probe. Samples (3 mm NMR tubes) were prepared in Kpi buffer 10 mM K2HPO4/KH2PO4; 50 mM KCl, at pH 6.8. D2O (8 %). The water signal was suppressed using excitation sculpting with gradients (zgesgppe; d1=2sec; 512 scans; time domain=64k). NMR data sets were treated using TopSpin 4.1.

### Supplementary Table

**Table S1.** Oligonucleotides sequences with a summary of the mass spectrometry data including their monoisotopic calculated and experimental mass values.

| name | sequence | Calculated Mass | Experimental Mass | ΔM (ppm) |
|---|---|---|---|---|
| control | 5'-**ketal**-TGT AGT AGG TTG TGT CTG G -3' | 6793.040 | 6793.038 | 0.3 |
| G4-2 | 5'- TGG GAG -3' | 1951.241 | 1951.183 | 29.7 |
| Lipid G4-2 | 5'-**ketal**-TGG GAG -3' | 2705.713 | 2705.631 | 30.3 |
| G4-3 | 5'- TGG GAG T -3' | 2271.264 | 2271.193 | 31.3 |
| Lipid G4-3 | 5'-**ketal**- TGG GAG T -3' | 3025.736 | 3025.646 | 29.7 |
| Lipid KRAS 21R | 5'-**ketal**- AGG GCG GTG TGG GAA GAG GGA-3' | 7416.641 | 7416.442 | 26.8 |

### Supplementary Figures

**Figure S1. Mass spectra of oligonucleotides.**
(**A**) Control, (**B**) G4-2, (**C**) Lipid G4-2, (**D**) G4-3, (**E**) Lipid G4-3, (**F**) Lipid KRAS 21 R

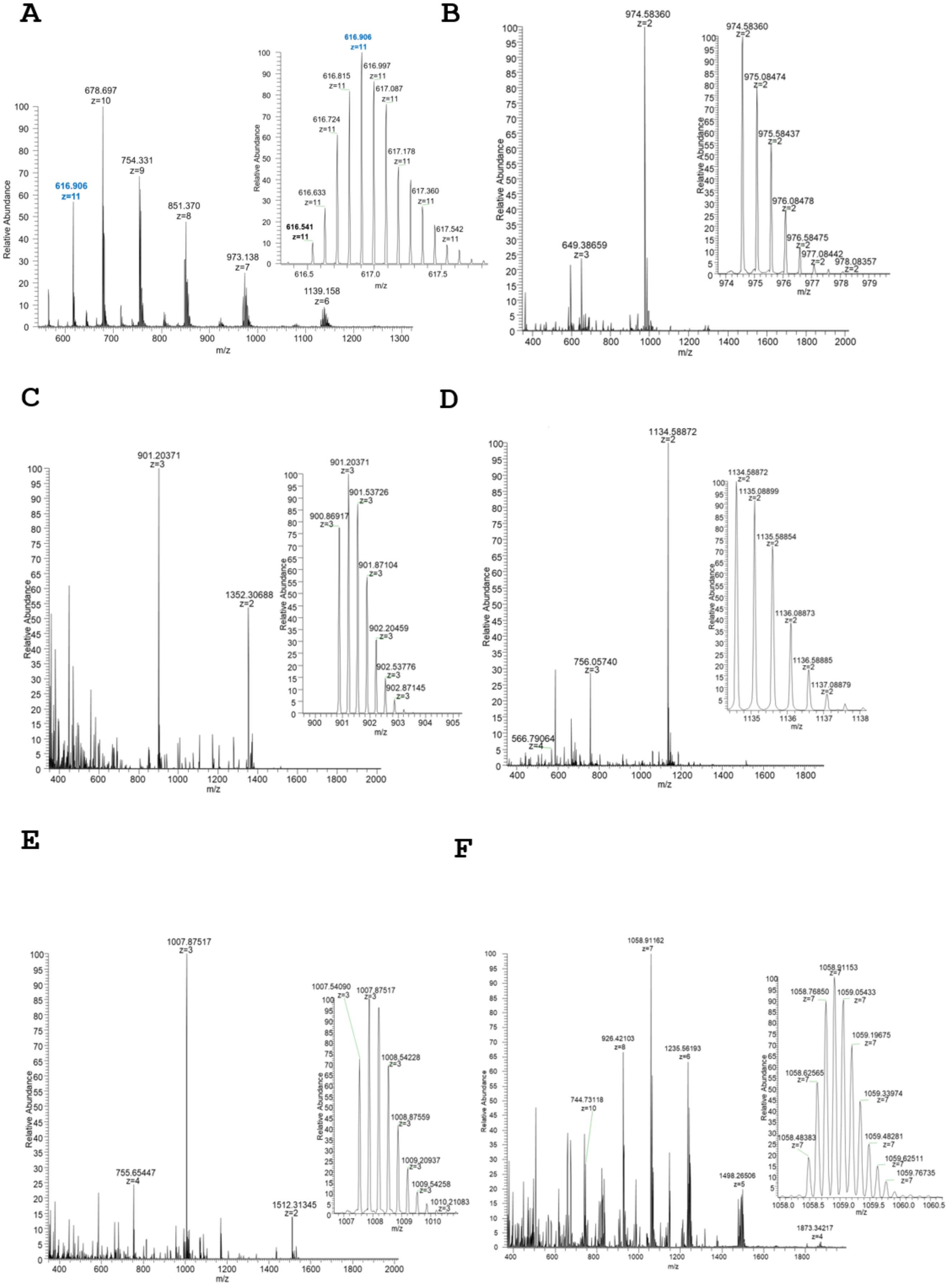

**Figure S2. HPLC chromatograms of oligonucleotides after purification.**
(**A**) Control, (**B**) G4-2, (**C**) G4-3, (**D**) Lipid G4-2, (**E**) Lipid G4-3 and (**F**) Lipid KRAS 21R

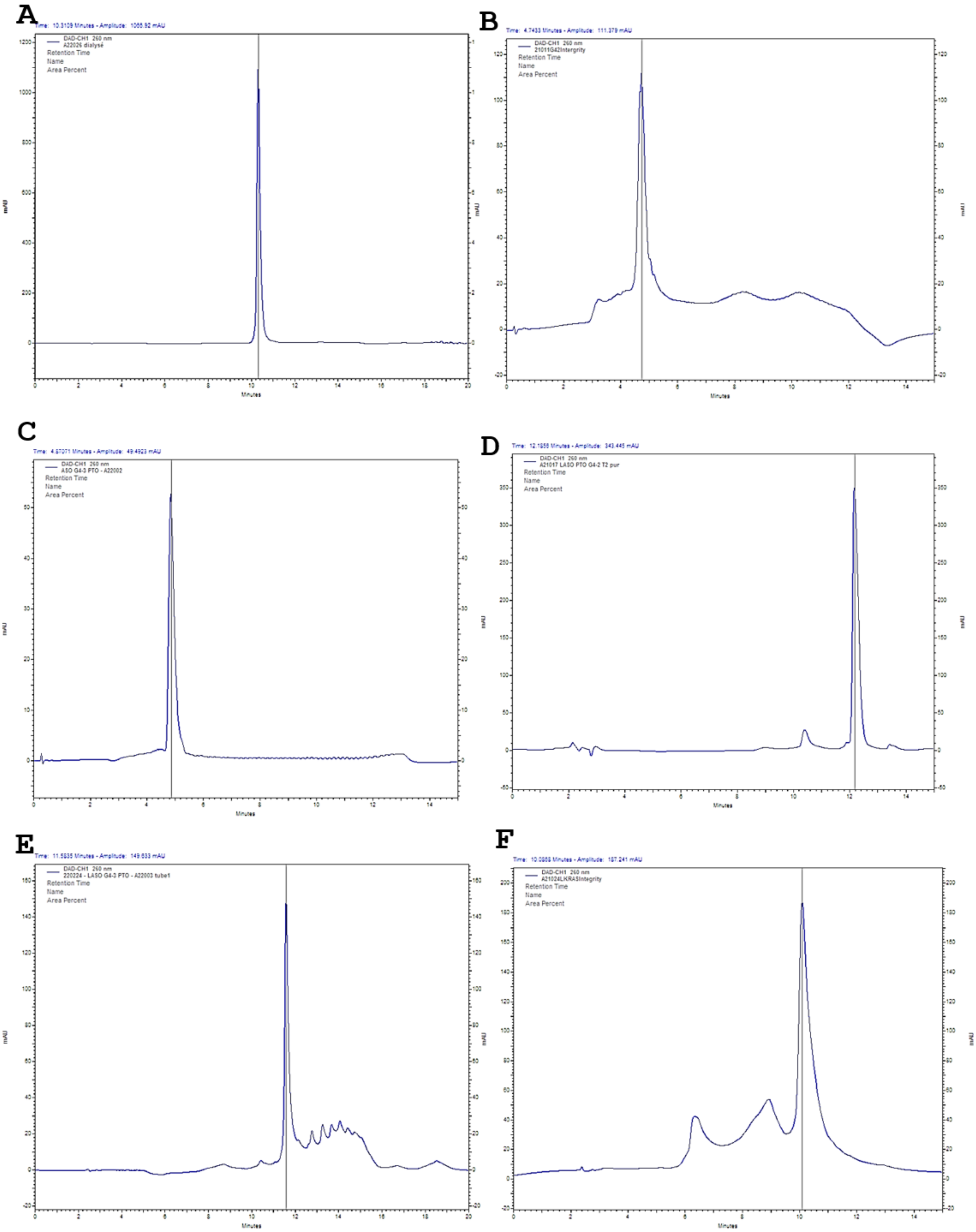

**Figure S3. DLS determined nano-object formation of lipid-conjugated oligonucleotides in physiological saline conditions.**

Blue and red graphs represent duplicates. (**A**) Lipid conjugated G4-2 oligonucleotides assemble into supramolecular structures that have a diameter of 223.9 nm and 26.02nm in intracellular (upper graph) and extracellular conditions respectively.

(**B**) Lipid G4-3 oligonucleotides form nano-objects with a diameter of 13.37 nm and 15.44 nm. (**C**) The native G quadruplex mimicking the KRAS promoter sequence conjugated to lipid forms nanostructures with a diameter of approximately 20nm in both intracellular and extracellular buffers. (**D**) Agarose gel showing the resolved nanobjects resulting from self-assembly of NLG4s

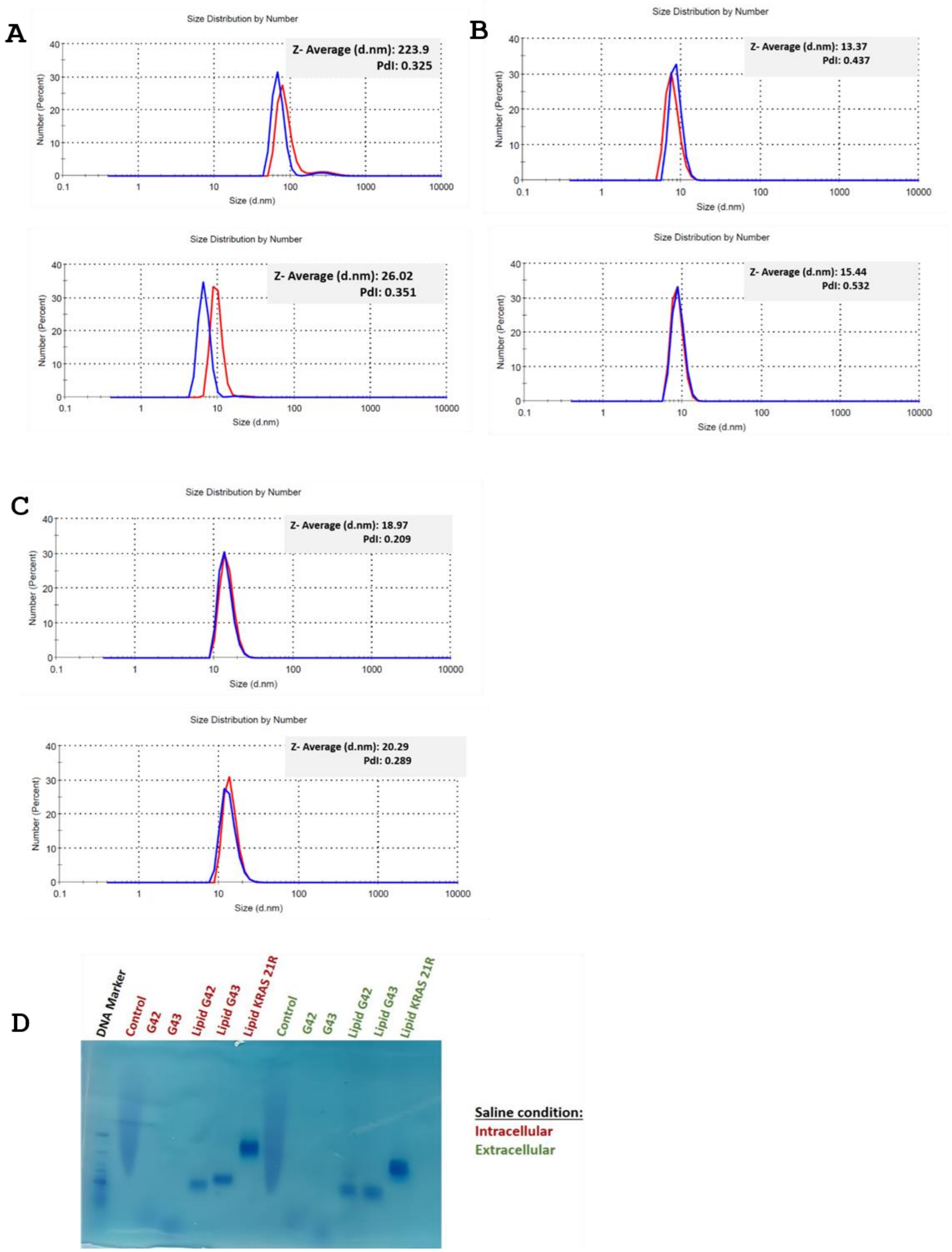

**Figure S4. Characterization of G4 formation and interaction with a G quadruplex binding protein.**

(**A**) Determination of the fluorescence threshold for G4 formation using Thioflavin T assay. (**B**) CD melting experiment, from 5 to 90°C, of G-quadruplexes under intracellular and extracellular saline conditions. Lipid-conjugated sequences are more stable in comparison to the unmodified sequences G4-2 and G4-3. Of the lipid-modified sequences, tetramolecular G quadruplexes tend to be more stable than the KRAS native sequence**.** (**C**) Electrophoretic mobility shift assay. Oligonucleotides are resolved in the absence (-) and presence (+) of GST tagged UP1. G-quadruplexes interact with the protein hence retarded on the gel. The G4 together with their protein interaction are visualized with the help of Thioflavin T fluorescence enhancement.

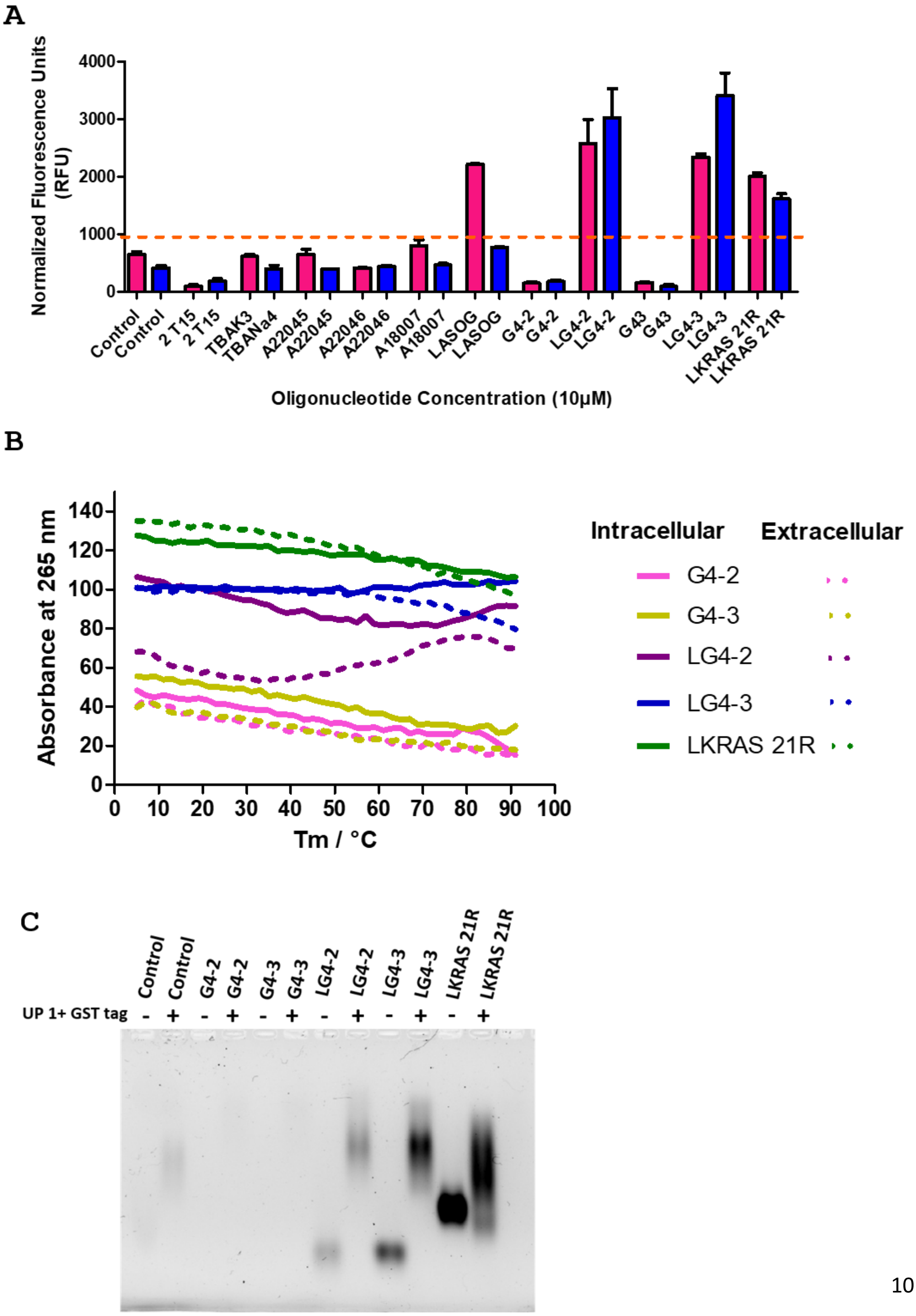

**Figure S5. Dose-dependent inhibitory effect of G4 lipid-conjugated oligonucleotides on PDAC cell survival at 1, 5, and 10μM.**

HPAFII cell viability was evaluated 72h after G4 oligonucleotides +/- lipids treatments (NT= Non Treated cells). All the experiments were performed in independent triplicate. Statistically significant differences between the treatments were determined by analysis of variance (one-way ANOVA). p-values of Tukey's post-hoc analyses are presented. *P<0.05, **P<0.01 and ***P<0.001.

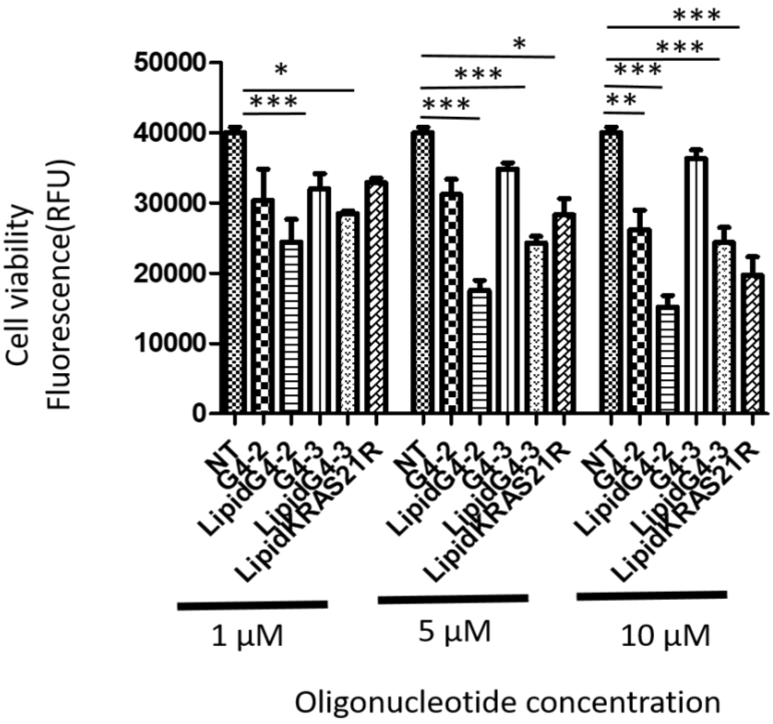

**Figure S6. NLG4-based treatments induce the reduction of BxPC3 PDAC cell survival, migration and growth with no effect on normal cells.**

A and B- PDAC BxPC3 cell line (**A**) and normal human bronchial epithelium BEAS-2B cell line (**B**) survival after the administration of lipid-modified G4 oligonucleotides and controls (NT = Non Treated and CTRL= scrambled oligonucleotide) at the concentration of 10µM for 72h. (**C**) BxPC3 cells migration capacities were evaluated using transwells, 72h after indicated treatments. Representative images of migrated cells stained with crystal violet are showed in the first line (n=3 transwells per treatment). For quantification, crystal violet was solubilized in ethanol and the absorbance at 555 nm was measured. (**D**) Spheroids diameters were measured 2 weeks after pre-treated BxPC3 cells were embedded in Matrigel™, using the Optika PROView analysis software. Representative images of the spheroids are showed in the first line (scale bar = 100 µm). The quantification was performed on n=6/7 spheroids per conditions. (**E**) A viability assay was performed on BxPC3 cells treated with G4 oligonucleotides and controls (NT and CTRL) at 10 µM in combination (+GEM) or not (-GEM) with gemcitabine (GEM) chemotherapy at 5 µM for 3 days. (**F**) GEM-resistant BxPC3 cell line was treated with G4 oligonucleotide decoys and control conditions at 10 µM for 72h and the viability of the cells was measured using CellTiter Blue reagent. Statistically significant differences between the treatments were determined by analysis of variance (one-way ANOVA). p-values of Tukey's post-hoc analyses are presented. *P<0.05, **P<0.01 and ***P<0.001.

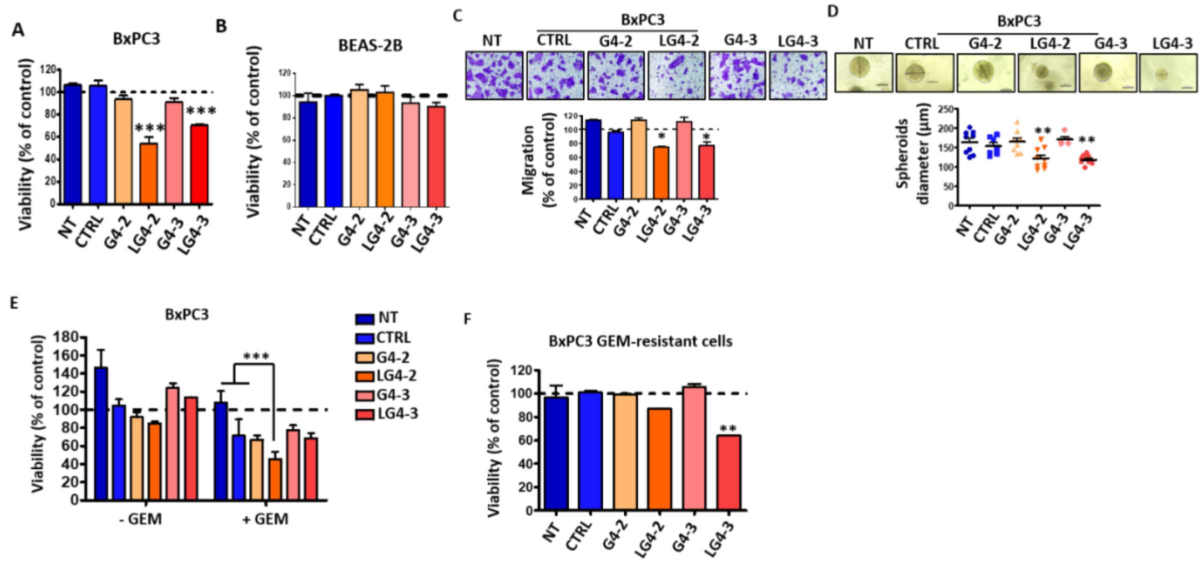
